# Targeting Tumor-derived Sphingosine Kinase 2 Unleashes Antitumor Immunity and Improves Survival of Mice with Group 3 Medulloblastoma

**DOI:** 10.64898/2026.08.12.744521

**Authors:** Sampurna Chatterjee, Pragya Kumar, Ashwin S. Kumar, Pin-Ji Lei, Meenal Datta, Yuhui Zhao, William W. Ho, Nilesh P. Talele, Patrik Andersson, Mark Duquette, Shuji Kitahara, Landry Blanc, Samantha J. Wong, Wilhelmus J. Kwanten, David H. Ebb, Torunn I. Yock, Veronique Dartois, Dai Fukumura, Dan G. Duda, Lei Xu, Hye-Jung Kim, Rakesh K. Jain

**Affiliations:** Edwin L. Steele Laboratories for Tumor Biology, Department of Radiation Oncology, Massachusetts General Hospital, Boston, MA 02114, USA; Harvard-MIT Division of Health Sciences and Technology, Massachusetts Institute of Technology, Cambridge, MA 02139, USA; Public Health Research Institute, New Jersey Medical School, Rutgers, The State University of New Jersey, Newark, New Jersey 07103, USA. Veronique Dartois Center for Discovery and Innovation, Hackensack Meridian Health, Nutley, NJ 07110, USA; Department of Cell Biology, Harvard Medical School, Boston, MA 02115, USA; Department of Pediatric Hematology/Oncology, Mass General Cancer Center, Massachusetts General Hospital, Boston, MA 02114, USA; Francis H. Burr Proton Therapy Center, Department of Radiation Oncology, Massachusetts General Hospital, Boston, MA 02114, USA; Department of Immunology, Dana Farber Cancer Institute, Harvard Medical School, Boston, MA 02115, USA; Immunology R&D, Translational Medicine & Bioinformatics, AstraZeneca, Boston, MA 02210, USA; Department of Tumor Microenvironment and Metastasis, H Lee Moffitt Cancer Center and Research Institute, Tampa, FL, USA; Cancer Biology PhD Program, University of South Florida, Tampa, FL, USA; Process Accelerator for Cell Therapy Manufacturing, Bioprocessing Technology Institute, Agency for Science Technology and Research, Singapore 138668, Singapore; Leman Biotech, Shenzhen, Guangdong, China; Department of Aerospace and Mechanical Engineering, University of Notre Dame, Notre Dame, IN 46556, USA; Department of Neurology at the Affiliated Hospital of Guangdong Medical University in Zhanjiang, Guangdong, China; Singapore Immunology Network (SIgN), Singapore 138648, Singapore; Department of Immunology, Trutino Biosciences, San Diego, CA 92121, USA; Faculty of Advanced Techno-Surgery, Institute of Advanced Biomedical Engineering and Science, Tokyo Women’s Medical University, Tokyo, 162-8666, Japan; University of Toulouse, CNRS, IPBS, Toulouse, France; Angelini Ventures Singapore, OUE Downtown, Singapore 068809, Singapore; Laboratory of Experimental Medicine and Paediatrics (LEMP) - Gastroenterology & Hepatology, University of Antwerp (UA), Belgium; Transplant Oncology and Therapeutics Program, Department of Surgery, Houston Methodist Academic Institute, Houston, TX 77030, USA; Department of Immunology Discovery, Genentech Inc., South San Francisco, California, USA

## Abstract

Group 3 medulloblastomas (G3MB) carry the worst prognosis among medulloblastoma subtypes, yet molecularly targeted therapies remain elusive. Standard treatments cause severe long-term morbidity in survivors. Here, we identify tumor-derived sphingosine kinase 2 (SPHK2) as an essential driver of G3MB initiation and progression. SPHK2 exacerbates local immunosuppression by suppressing cytotoxic T-cell and NK-cell activity while promoting regulatory T-cell infiltration. Genetic or pharmacologic SPHK2 inhibition using Opaganib attenuates pro-survival tumor signaling and restores anti-tumor immunity, significantly improving survival in syngeneic G3MB mouse models. Combining Opaganib with fractionated low-dose radiation (f-LDRT) further enhances antigen presentation and reprograms tumor-associated myeloid cells toward an anti-tumor phenotype. This combination therapy markedly prolongs survival without inducing significant toxicity. Overall, our study establishes SPHK2 as a previously unrecognized therapeutic target and presents a safe, effective, microenvironment-reprogramming regimen for G3MB.

**One Sentence Summary:** Direct inhibition of tumor-derived SPHK2 overcomes local immunosuppression and downregulates pro-survival signaling in Group 3 medulloblastoma, while combination with fractionated low-dose radiation further enhances anti-tumor immunity and significantly improves survival.

## Introduction

Medulloblastoma (MB) is the most common malignant pediatric brain tumor (Juraschka & Taylor, 2019). Genomic classification and risk stratification reveal MB to be a heterogenous disease with four distinct subgroups—WNT-driven, SHH-driven, Group 3, and Group 4—each with potential vulnerabilities for targeted therapy. Group 3 MB (G3MB) has the worst prognosis amongst the four subgroups with less than 50% of pediatric patients surviving up to 5 years (Menyhárt et al., 2019). G3MB is often associated with adverse prognostic factors such as large cell anaplastic (LCA) histology and *c-Myc* amplification, a master regulator of the tumor microenvironment (TME) (Cavalli et al., 2017; Juraschka & Taylor, 2019; D. Kumar et al., 2025; Lindsey et al., 2016; Tao et al., 2019). In contrast to targetable WNT and SHH signaling pathways that drive those MB subgroups, MYC protein is a poor therapeutic target in G3MB due to its pleiotropic nature (Hurlin, 2013; Miller et al., 2012). The TME is recognized as a key contributor to tumor progression and immunosuppression in G3MB (Pham et al., 2016; Snuderl et al., 2013). Thus, targeted therapies that can reprogram the immunosuppressive TME into an immunostimulatory one, have the potential to improve G3MB treatment and are desperately needed.

The sphingolipid signaling pathway, which includes bioactive sphingolipids such as ceramide, ceramide-1-phosphate (C1P), sphingosine, sphingosine-1-phosphate (S1P), and sphingosine kinases (SPHKs), is activated in neurons, oligodendrocytes, and endothelial cells (Mei et al., 2023; Nagahashi et al., 2012). S1P binds to S1P receptors (S1PR1-5), regulating multiple downstream signaling pathways (Maceyka & Spiegel, 2014; Vestri et al., 2017) to maintain brain development, cellular interaction, and metabolism. SPHKs phosphorylate sphingosine to S1P. Two SPHK isoforms—SPHK1 and SPHK2—have been reported with overlapping functions. Simultaneous *Sphk1/Sphk2* knockout (KO) in mice causes embryonic lethality due to vascular and neurological defects, whereas *Sphk1* KO and *Sphk2* KO mice develop normally (Mizugishi et al., 2005). SPHK1 aids tumor progression in breast, prostate, ovarian cancer, melanoma and glioblastoma (GBM) (Acharya et al., 2019; Chakraborty et al., 2019; Hart et al., 2019; C.- F. Lee et al., 2019; Young et al., 2009). Although SPHK2 is expressed in the cerebellum, little is known about the role of SPHK2 in pediatric central nervous system (CNS) tumors including G3MB.

Given that SPHK2 is dispensable during cerebellar development, we aimed to explore the mechanistic role of SPHK2 and its potential as a therapeutic target in G3MB using both genetic and pharmacological approaches. We chose Opaganib as a potent and selective inhibitor of SPHK2 in G3MB. This inhibitor binds to the sphingosine-binding site of SPHK2, with minimal off-target effects on other protein kinases and no reported systemic toxicity in murine models (Britten et al., 2017; French et al., 2010). A phase 2 trial (NCT04207255) is currently underway investigating Opaganib’s ability to overcome resistance to androgen receptor pathway inhibitors in metastatic castration-resistant prostate cancer (mCRPC) patients (Kucuk et al., 2021).

Current standard of care (SOC) for children (3 years or older) with MBs involves craniospinal irradiation (CSI) with a boost to the original tumor bed (Juraschka & Taylor, 2019; V. Kumar et al., 2017; Kuzan-Fischer et al., 2018; Perek-Polnik et al., 2023). CSI has been shown to induce immunosuppression in newly diagnosed MB patients by reducing circulating levels of total lymphocytes and increasing immunosuppressive regulatory T cells (Gururangan et al., 2017). In pre-clinical studies, anti-tumor response to radiation therapy depends on the dose per fraction, the number of fractions, and the total dose (Demaria et al., 2015; Demaria & Formenti, 2012; Y. Lee et al., 2009; Lugade et al., 2005). In fact, fractionated low dose radiation (f-LDRT) can stimulate the production of tumor-associated antigens (TAAs) and induce MHC class I and II molecules in various cancer cell lines (lung, colon, prostate, and MB) (Arnold et al., 2018; Das et al., 2017; Minturn et al., 2021). Therefore, f-LDRT seems to be an attractive strategy to induce immune stimulation and reprogram the TME in combination with Opaganib in preclinical models of G3MB.

Here, we show that tumor-derived SPHK2 signaling promotes tumor progression and immunosuppression in G3MB. Moreover, selectively inhibiting SPHK2 with Opaganib enhances anti-tumor immunity and nearly doubles the survival of mice bearing syngeneic models of G3MB with no symptoms of overt toxicity.

## Results

### Sphingosine kinase 2 and downstream signaling molecules are overexpressed in human and murine G3MB

We detected abundant SPHK2 in both human and murine MB samples using immunohistochemistry (IHC). Using an MB tissue array, we examined SPHK2 in seven pediatric MB patient tumor tissues (**Figure 1A,B)** that displayed large-cell anaplastic (LCA) histology (**Figure S1A);** LCA histology is frequently associated with G3MB (Louis et al., 2016; Northcott et al., 2012; Pei et al., 2012). SPHK2 scoring was significantly higher in the human MB tissues, compared to normal cerebellum (**Figure 1).** We also detected abundant SPHK2 in two orthotopic murine G3MB models, hereafter termed MYC1 (Kawauchi et al., 2012) and MYC2 (Pei et al., 2012) (**Figure 1C** and **Figure S1B- D)**. Murine *Sphk2* mRNA expression relative to normal cerebellum was higher than the relative mRNA levels of its isoform, *Sphk1,* in both MYC1 and MYC2 tumors (**Figure 1D**). Murine mRNA expression levels of *S1pr1* and *S1pr2*, receptors of SPHK1/2 were significantly elevated in MYC1 and MYC2 tumors compared to normal cerebellum (**Figure 1E**). Moreover, we found more cells positive for pSTAT3, which is downstream of S1PRs (Nguyen, 2014), at Ser727 in MYC1 tumors compared to normal cerebellum (**Figure 1F**). Taken together, our data show in G3MB that SPHK2 and its receptors *S1pr1* and *2* are highly expressed and STAT3, a downstream molecule in the sphingolipid signaling pathway, is phosphorylated.

**Figure 1.**
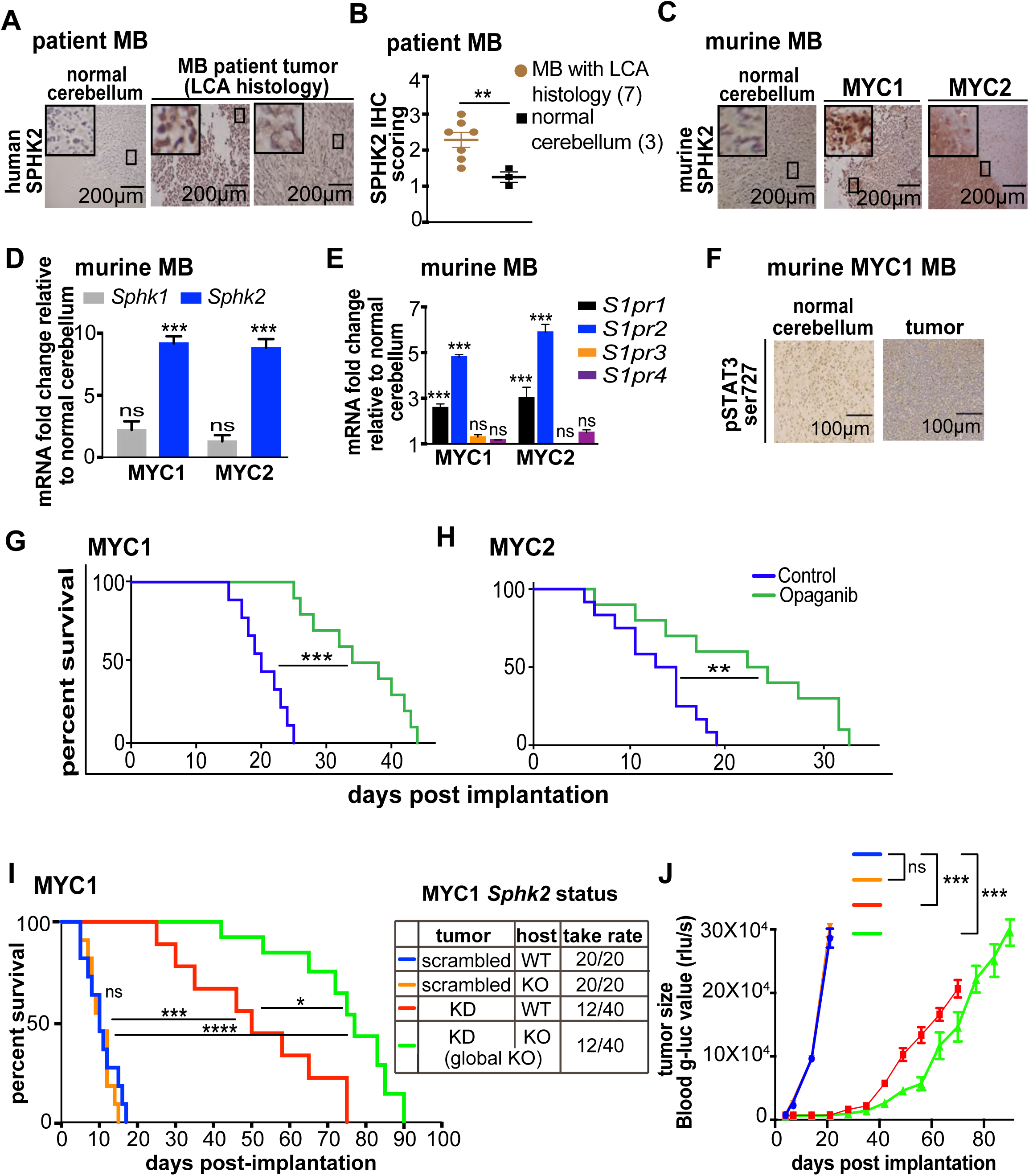
Sphingosine kinase 2 (SPHK2) is overexpressed in G3MB, and its inhibition enhances survival in G3MB models. **(A)** IHC images of MB patient cerebellum tissues with LCA histology stained for SPHK2. **(B)** SPHK2-positive histological score in pediatric patient MB tumors (n=7) and normal cerebellum (n=3). Histological score was assigned based on percentage of tumor cells with positive stain as <20%=0, 20-50%=1, 50-80%=2 and >80%=3. ** = p≤0.01. Error bars represent SEM. **(C)** IHC images of SPHK2 staining in murine MYC1 and MYC2 G3MB and normal cerebellum. **(D)** qPCR of *Sphk1* and *Sphk2,* in murine MYC1 and MYC2 tumors, normalized to normal cerebellum. *** = p≤0.001, n=6/group. Error bars represent SEM. **(E)** qPCR of *S1pr1*, *S1pr2, S1pr3* and *S1pr4* in MYC1 and MYC2 G3MB, normalized to normal cerebellum. *** = p≤0.001, n=2/group. **(F)** IHC images pSTAT3 at Ser727 staining in murine MYC1 MB and normal cerebellum. **(G, H)** Survival curves in MYC 1 (**G**) and MYC2 (**H**) models with indicated treatments, comparing median survival. Opaganib: 200mg/kg daily oral gavage. n=15/group. **p≤0.01, ***p≤0.001. **(I)** Survival curve in MYC 1 model in Sphk2 WT, KD or KO in host vs tumor as indicated, comparing median survival. * = p≤0.05, *** = p≤0.001, **** = p≤0.0001, n=12/group. **(J)** Tumor burden measured by Gaussia luciferase (g-luc) signal in blood in Sphk2 WT, KD or KO in host vs tumor as indicated. ***=p≤0.001. Error bars represent SEM.

### SPHK2 inhibition delays tumor growth and doubles the survival of mice bearing G3MB

To examine the impact of SPHK2 inhibition on G3MB progression, we orthotopically implanted murine MYC1 and MYC2 tumor cells into the cerebellum of immunocompetent C57Bl/6 mice. Mice started developing tumors within 1 week from implantation -- monitored by ultrasound, circulating g-luc values or whole-body imaging (**Figures S1B-D**) as previously established (Askoxylakis et al., 2017; Snuderl et al., 2013). We treated mice with established tumors with Opaganib (Selleckchem), a small-molecule inhibitor of SPHK2. Opaganib significantly improved median survival by 16 days in MYC1 (by 80 %, p<0.001) and 10 days in MYC2 (by 76.9%, p≤0.01) (**Figures 1G, H)** and inhibited tumor growth in MYC1 (by 80%, p<0.001) and MYC2 (by 77%, p<0.01) G3MB models (**Figure S1E**). Consistent with its low-toxicity profile (Britten et al., 2017; French et al., 2010), Opaganib-treated mice showed no clinical symptoms of toxicity, such as hunched posture or hair loss, and lost less than 15% body weight compared to untreated control mice in both models (**Figure S1F**). Next, to test if Opaganib could reach the brain tumors, we performed mass spectrometry (MS) of tumors from untreated mice and mice treated with Opaganib. MS data showed increased Opaganib concentration in MB tumors in treated MYC1 (∼0.5µM) and MYC2 (∼0.7µM) mice compared to untreated controls (<0.1µM) **(Figure S1G)**. Opaganib also reduced viability in MYC1 MB cells at 0.7µM and 1µM concentrations (**Figure S1H**).

### Tumor-derived SPHK2 mediates G3MB development and progression

Since SPHK2 is expressed in diverse CNS and stromal cells, such as astrocytes, glial cells, and neurons, we next analyzed the role of sphingosine signaling in tumor versus stromal cells in driving MB tumorigenesis and progression in mice. MYC1 MB cells harboring shRNA1 and 2 to knockdown (KD) *Sphk2* (efficiency: 60-80%) (**Figure S1I**) were orthotopically implanted in wild-type (WT) or *Sphk2* knockout (KO) mice. We then compared tumor development and survival amongst the following groups: 1) scrambled shRNA MB cells in *Sphk2* WT mice (control group), 2) scrambled shRNA MB cells in syngeneic *Sphk2*-KO mice, 3) *Sphk2*-KD MB cells in *Sphk2* WT mice, and 4) *Sphk2*-KD MB in syngeneic *Sphk2*-KO mice (*Sphk2* global KO). We found a lower tumor take rate in mice with *Sphk2-*KD MBs, both in WT mice and *Sphk2*-KO mice (∼30% in both groups) compared to control or *Sphk2*-KO mice bearing scrambled shRNA MB cells (100% in both groups) (**Figure 1I**). Both*Sphk2-*KD MB groups had a significantly longer median survival than scrambled MB groups. Median survival in *Sphk2* KD MB cells in *Sphk2* WT mice was 50 days (p<0.001), and the longest survival was seen in *Sphk2* KO mice with *Sphk2* KD MBs with a median survival of 77 days (p<0.0001) (**Figure 1I**). In contrast, there was no difference in survival between scrambled shRNA MB-bearing WT mice and *Sphk2*-KO mice (median survival 10 days in both cases) (**Figure 1I**). In these mice, the tumor take rate was 100%, and they all exhibited rapid progression compared to both*Sphk2-*KD groups (**Figure 1J**). These data indicate that tumor-derived SPHK2 mediates G3MB engraftment and progression. However, in its absence, host-derived SPHK2 can aid tumor progression in G3MBs. Targeting sphingosine kinase 2 genetically or pharmacologically significantly improves survival in mice bearing G3MB tumors.

### SPHK2 inhibition disrupts tumor-promoting signaling and unleashes an immune response against G3MB

We investigated whether inhibiting SPHK2 could disrupt the dysregulated SPHK2/S1PR cascade at transcriptional and post-transcriptional levels. Western blot analysis of the downstream signaling revealed that Opaganib decreased S1PR2 in MYC1 tumors *in vivo* and the MYC1 cell line *in vitro* (**Figures 2A,B**), without affecting S1PR1 levels (**Figure 2A**). Opaganib also reduced phosphorylation of STAT3 at Ser727 in MYC1 tumors and the MYC1 cell line *in vitro* (**Figures 2A,B**). pERK was consistently reduced in MYC1 cell line under Opaganib in a dose-dependent manner (**Figure 2C**). Strikingly, Opaganib reduced cMYC levels -- a pivotal driver of G3MB tumorigenesis -- in the MYC1 cell line (**Figure 2C**). To gain further mechanistic insights into the role of targeting SPHK2 in improving survival, we performed RNA sequencing on *Sphk2* WT tumors (scrambled shRNA MB cells in *Sphk2* WT mice), and *Sphk2* global KO tumors (*Sphk2*-KD MB cells in *Sphk2*-KO mice). We identified multiple differentially expressed genes (DEGs) associated with MB tumorigenesis that were altered in *Sphk2* global KO tumors. *Cdkn2a*, *Bid,* and *Casp3* were upregulated in *Sphk2* global KO tumors compared to *Sphk2* WT (**Figure 2D**). *E2f*, the principal target of the tumor suppressor pRB (Chen et al., 2009), and *Mad2l2*, a component of the mitotic spindle assembly checkpoint and a good prognostic marker in colorectal cancer (Li et al., 2018), were upregulated in MYC1 *Sphk2* global KO tumors, while *Fgfr3* was downregulated compared to Sphk2 WT (**Figure 2D**). Concurrently, gene set enrichment analysis (GSEA) showed significant upregulation of gene sets associated with cell cycle checkpoint and arrest, transcriptional regulation by TP53, immune response such as TCR signaling, and antiviral mechanism by IFN-stimulated genes in the MYC1 *Sphk2* global KO tumors compared to *Sphk2* WT (**Figure 2E**, **Supplemental Tables 1-3)**. Collectively, these results indicate that targeting SPHK2 in G3MB not only disrupts downstream tumor-promoting signaling but also activates the immune response in the TME of G3MB.

**Figure 2.**
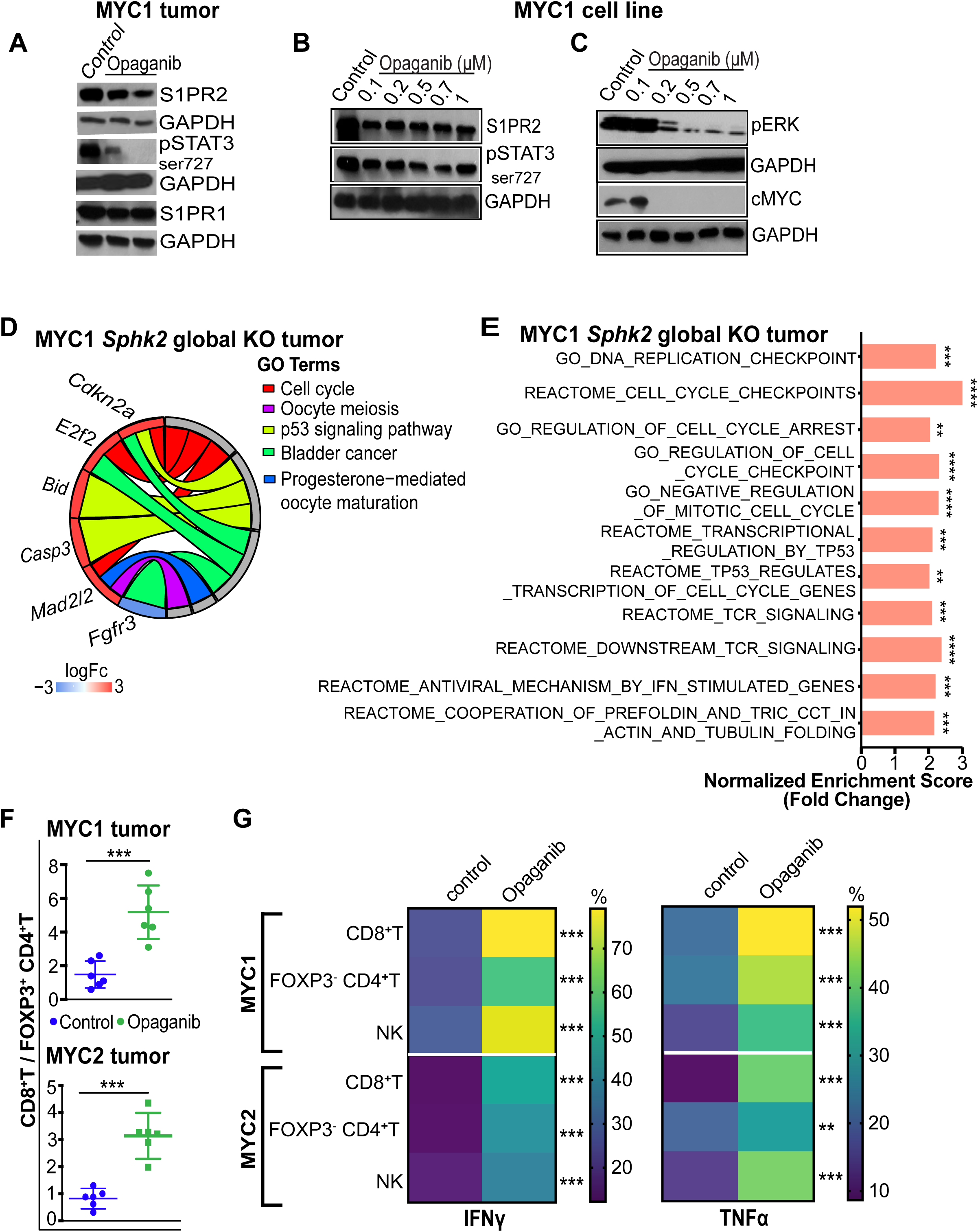
SPHK2 inhibition disrupts tumor-promoting signaling and unleashes CD8+, CD4+, and natural killer (NK) lymphocyte activity in G3MB tumors. **(A)** WB showing levels of indicated proteins in MYC1 tumors from mice treated with Opaganib. Opaganib: 200mg/kg daily oral gavage. **(B, C)** WB showing levels of indicated proteins in MYC1 cell line treated with Opaganib at indicated concentrations for 72 hours. **(D)** GO pathway enrichment analysis of bulk RNA sequencing of MYC1 tumors. Differential gene expression was obtained using edgeR package (version 3.26.8). LogFc≥|2|, FDR q-value<0.05. The complete lists of differentially expressed genes for *Sphk2* global KO tumors versus *Sphk2* MYC1 WT tumors are in the **Supplemental Tables 1** and 2. **(E)** Bulk RNA sequencing GSEA of MYC1 tumors. ** = FDR q-value≤0.01, *** = FDR q- value≤0.001, ****FDR q-value≤0.0001. The complete GSEA report of *Sphk2* global KO tumors and MYC1 *Sphk2* WT (scrambled) tumors is in **Supplemental Table 3**. **(F)** Flow cytometry of MYC1 and MYC2 tumors treated with Opaganib showing ratio of CD8^+^T (CD45^+^CD8^+^) cells to FOXP3^+^ CD4^+^T (CD45^+^CD4^+^) Tregs. Opaganib: 200mg/kg daily oral gavage *** = p≤0.001, n=6/group. Error bars represent SEM. **(G)** Heatmap showing flow cytometry analysis of Opaganib treated MYC1 and MYC2 tumors for indicated populations. ** = p≤0.01, *** = p≤0.001 n=6/group.

### Pharmacological inhibition of SPHK2 activates CD8^+^, CD4^+^, and natural killer (NK) cells in G3MB tumors

Since our RNA sequencing analysis indicated activation of the immune response via the TCR pathway in *Sphk2* global KO tumors, we next asked whether pharmacological inhibition of SPHK2 could reprogram the T, NK, and NKT cells in G3MB. The ratio of CD8^+^T cells to FOXP3^+^CD4^+^ T regulatory cells (Tregs) increased significantly in Opaganib-treated tumors compared to controls in both MYC1 and MYC2 tumors (**Figure 2F**). Opaganib significantly increased the percentage of CD8^+^T cells, FOXP3^-^CD4^+^T conventional cells (Tcon), and NK cells secreting inflammatory cytokines such as IFNγ and TNFα compared to controls in both models (**Figure 2G)**. Combined CD8^+^T and NK cell depletion in conjunction with Opaganib significantly abrogated the survival benefit compared to Opaganib-treated mice alone or in conjunction with either CD8^+^T or NK cell depletion (**Figure S2A**). These results indicate that CD8+ T- and NK cells contribute to the survival benefit from Opaganib.

### SPHK2 inhibition facilitates cross-presentation of glycolipid antigens to NKT cells in a CD1d-restricted manner

Since MBs are known to lack surface expression of MHC-I or components of MHC-I processing machinery, we investigated alternative mechanisms of T and NK cell activation in the tumor under SPHK2 inhibition (Kurdi et al., 2023; Smith et al., 2009). We used flow cytometry to analyze CD1d, a non-classical MHC class I-like glycoprotein capable of presenting glycolipids and phospholipids to Natural Killer T (NKT) cells. We discovered that both MYC1 and MYC2 expressed CD1d on the cell surface (**Figure 3A**). Presentation of α-Galactosyl Ceramide (α-GalCer), a CD1d binding antigen, on CD1d tetramers significantly increased the frequency of antigen-specific invariant natural killer (iNKT) cells under Opaganib in either tumor model compared to control (**Figure 3B**). Next, to evaluate the tumor-killing capacity of these iNKT cells, we performed an in vitro cytotoxicity assay. iNKT cells sorted from treated tumors were significantly cytotoxic towards CD1d^+^ MYC1 cells pre-pulsed with α-GalCer, compared to vehicle control (**Figure 3C**). Importantly, these iNKT cells in the treated group produced high levels of IFNγ, TNFα, IL-4, and IL-12, cytokines in the supernatant that are commonly associated with cytotoxic activity of T, NK, and NKT cells (**Figure 3D**). Tumor lysates and peripheral blood from Opaganib-treated mice showed a similar signature of upregulated cytokine levels compared to controls (**Figures 3E,F)**. These results indicate that MB cells can upregulate CD1d ligand upon SPHK2 inhibition and trigger robust CD1d-restricted iNKT cell cytotoxicity.

**Figure 3.**
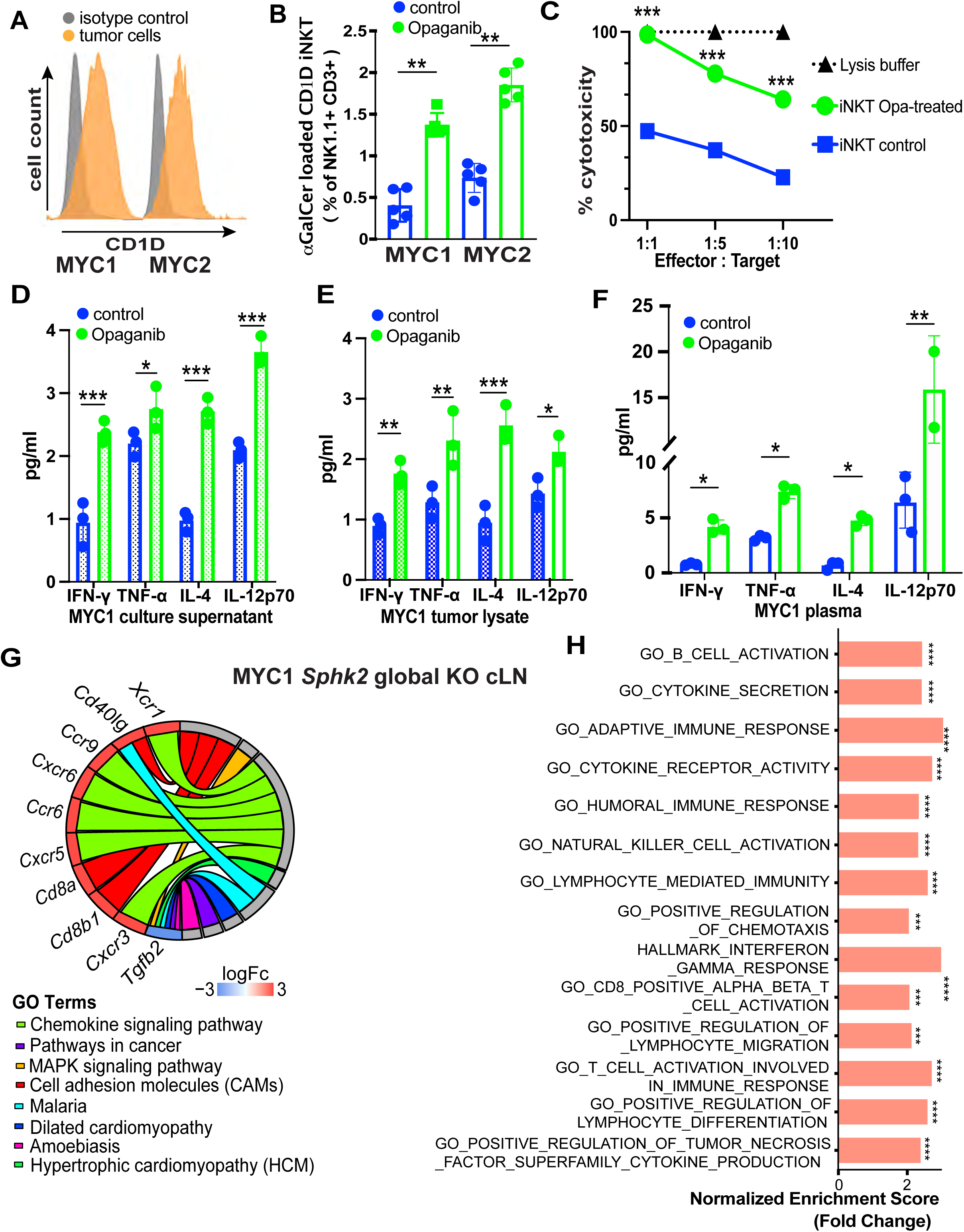
SPHK2 inhibition enhances anti-tumor immunity in the draining lymph nodes. **(A)** Flow cytometry of MYC1 and MYC2 tumors. Representative image of mean fluorescence intensity (MFI) of CD1d are shown. **(B)** Flow cytometry of MYC1 and MYC2 tumors treated with Opaganib showing percentages of iNKT (NK1.1^+^CD3^+^) cells are shown. n=4-5/group. * = p≤0.05, *** = p≤0.001. Error bars represent SEM. **(C)** *In vitro* cytotoxicity assay showing tumor-killing capacity of iNKT cells sorted from MYC1 tumors against CD1d+ MYC1 cells pre-pulsed with α-GalCer at indicated effector:target ratios. iNKT cells from Opaganib-treated tumors are compared with iNKT cells from control tumors. *** = p≤0.001. **(D)** Cytokine levels in supernatants from the in vitro iNKT cell cytotoxicity assay showing IFNγ, TNFα, IL-4, and IL-12p70. n=2-3/group * = p≤0.05, ** = p≤0.01. Error bars represent SEM. **(E)** Cytokine analysis of MYC1 tumor lysates from mice treated with Opaganib or control showing IFNγ, TNFα, IL-4, and IL-12p70. n=2-3/group * = p≤0.05, ** = p≤0.01, *** = p≤0.001. Error bars represent SEM. **(F)** Cytokine analysis of plasma from mice bearing MYC1 tumors and treated with Opaganib or control showing IFNγ, TNFα, IL-4, and IL-12p70. n=2-3/group * = p≤0.05, ** = p≤0.01, *** = p≤0.001. Error bars represent SEM. **(G)** GO pathway enrichment analysis of bulk RNA sequencing of MYC1 draining cervical lymph nodes (cLN). Differential gene expression was obtained using edgeR package (version 3.26.8). LogFc≥|2|, FDR q-value<0.05. The complete lists of differentially expressed genes for *Sphk2* global KO cLNs versus *Sphk2* MYC1 WT cLNs are in the **Supplemental Tables 4** and 5. **(H)** Bulk RNA sequencing GSEA of MYC1 cLNs. ** = FDR q-value≤0.01, *** = FDR q-value≤0.001, ****FDR q-value≤0.0001. The complete GSEA report of *Sphk2* global KO cLNs and MYC1 *Sphk2* WT (scrambled) cLNs is in **Supplemental Table 6.**

Since draining lymph nodes (LNs) are critical for the initiation of tumor-specific immune responses (Murphy & Griffith, 2016), we next investigated whether the immune- modulatory effect of genetic and pharmacological inhibition of SPHK2 extended to the bilateral superficial cervical LNs (cLNs). Bulk RNA sequencing of cLNs from the *Sphk2* WT (scrambled) and *Sphk2* global KO mice identified key DEGs associated with immune response (n=4/group) with greater than 2-log fold change and an FDR cut-off value set as q-value <0.05. Immune-stimulatory molecule *CD40lg* and T lymphocyte markers *CD8a* and *CD8b1* were upregulated in *Sphk2* global KO cLNs compared to *Sphk2* WT cLNs (**Figure 3G**). Cytokines responsible for T and B cell development and tissue-specific homing, such as *Ccr6, Ccr9, Cxcr3, Cxcr5, Xcr1* and *Cxcr6*, were upregulated while *Tgfb2* was downregulated in the *Sphk2* global KO cLNs compared to *Sphk2* WT cLNs (**Figure 3G**). GSEA analysis revealed upregulation of multiple pathways associated with trafficking, differentiation, activation and cytotoxicity across lymphocyte populations such as T, NK, and B cells in the *Sphk2* global KO cLNs compared to *Sphk2* WT cLNs (**Figure 3H**, **Supplemental Tables 4- 6)**. The percentage of APC DC-like cells increased significantly in the treatment arm of MYC1 tumors but not in MYC2 tumors, indicating inter-tumoral heterogeneity (**Figure S2B**). In the cLNs, the percentage of APCs such as macrophages (Mφs) and DC-like cells co-expressing MHCI, MHCII, and CD86 remained unaltered under Opaganib compared to control in both models **(Figure S2B).** The varied degrees of DC infiltrations in tumors and lack of increased influx of other APCs in tumors and LNs, along with downregulation of antigen-receptor signaling in the LNs, indicate that G3MBs lack a crucial step in the activation of anti-tumor immune response, suggesting the need for additional therapy.

### Fractionated low-dose radiation (f-LDRT) potentiates anti-tumor immunity in *Sphk2* global knockout G3MB

Since Opaganib treatment did not decrease the fraction of TAMs or MGs, we hypothesized that adding f-LDRT would enhance the potential for tumor-associated antigen (TAA) presentation and expression based on recent studies (Arnold et al., 2018; Das et al., 2017). To this end, by testing multiple doses and schedules, we identified an optimum radiation dose of 10.8 Gy fractionated in 1.8 Gy fractions given every 3 days to the tumor area of the cerebellum bearing *Sphk2* WT tumors (scrambled shRNA MB cells in *Sphk2* WT mice). We observed the highest mRNA levels of *Cd1d,* an antigen-presenting molecule for NKT cells (D. Liu et al., 2013), in the 10.8 Gy-treated tumors compared to sham RT control as well as other f-LDRT doses **(Figure S2C)**. Mice in the 10.8Gy radiation group showed limited weight loss with some recovery over other radiation dose/schedules (**Figures S2D,E**). No hair loss or skin rashes were observed with this fractionated dose.

To understand the molecular mechanism of innate immune sensing of RT-induced genotoxicity, we next investigated whether f-LDRT induced damage-associated molecular patterns (DAMPs) in the TME. Cytosolic DNA concentrations in f-LDRT -treated MYC1 cells were aberrantly higher than in controls (**Figure 4A**). f-LDRT also induced activation of molecules of the pGAS/STING pathway, including pTBK1 and pIRF3 in cDC1 (**Figures 4B,C**) and upregulated MHCI expression on cDC1 (**Figure 4D**). Next, we tested whether our selected dose of f-LDRT in *Sphk2* WT G3MB or in combination with *Sphk2* global KO primes the immune response. Mean fluorescence intensity (MFI) of MHCII on cDC2 and TAMs increased only in tumors receiving f-LDRT compared to *Sphk2* WT, indicating that this is an f-LDRT-driven effect (**Figures 4E,F**). In the tumor-draining lymph nodes, the fractions of MHCII^+^CD86^+^ Mφs in f-LDRT treated cLNs were higher compared to *Sphk2* WT and *Sphk2* global KO arms (**Figure 4G**). The fraction of cDC2 increased significantly in all the f-LDRT treatment arms in cLNs compared to *Sphk2* WT (**Figure 4G**). This was associated with a significant increase in the fraction of MGs in the arms receiving f-LDRT in tumors (**Figure S3A**) but not TAMS or DCs in the tumors or Mφs or DCs in tumors or cLNs (**Figures S3B-E**). Overall CD45^+^ cell count in the tumors was significantly improved in the *Sphk2* global KO and the combination arm over control (**Figure S3F**).

**Figure 4.**
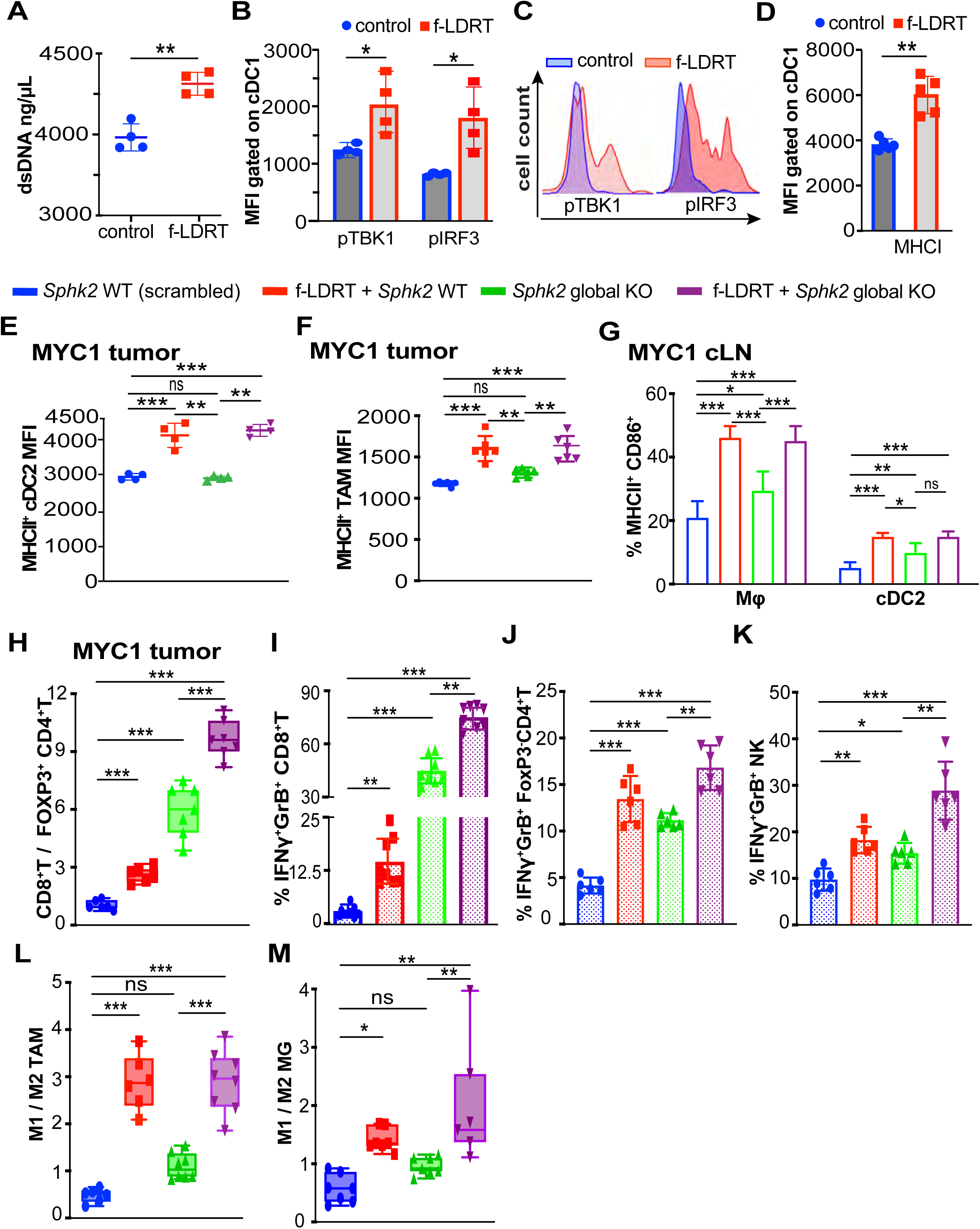
Fractionated low-dose radiation (f-LDRT) potentiates anti-tumor immunity in *Sphk2* global knockout G3MB. **(A)** Cytosolic double-stranded DNA (dsDNA) concentration in MYC1 cells treated with f- LDRT or control. n=4/group ** = p≤0.01. Error bars represent SEM. **(B)** Flow cytometry of cDC1 from MYC1 tumors treated with f-LDRT or control showing activation of the cGAS-STING pathway, as indicated by mean fluorescence intensity (MFI) of pTBK1 and pIRF3. n=4/group * = p≤0.05, ** = p≤0.01. Error bars represent SEM. **(C)** Flow cytometry of MYC1 tumor. Representative image of mean fluorescence intensity (MFI) of pTBK1 and pIRF3are shown. **(D)** Flow cytometry of cDC1 from MYC1 tumors treated with f-LDRT or control showing MHC I expression by mean fluorescence intensity (MFI). ** = p≤0.01. Error bars represent SEM. **(E, F)** Flow cytometry of MYC1 tumors under Sphk2 global KO vs WT, treated with fLDRT. Mean fluorescence intensity (MFI) of MHCII^+^ cDC2 **(E)** and MCHII^+^ TAM **(F)** are shown. n=4/group for **E**, n=6/group for **F**. ** = p≤0.01, *** = p≤0.001. Error bars represent SEM. **(G)** Flow cytometry of MYC1 cLNs under Sphk2 global KO vs WT, treated with fLDRT. Percentages of MHCII^+^CD68^+^ macrophages and cDC2 populations are shown. n=6/group. * = p≤0.05, *** = p≤0.001. Error bars represent SEM. **(H-M)** Flow cytometry of MYC1 tumors under Sphk2 global KO vs WT, treated with fLDRT for indicated populations. n=6-7/group for **H**, n=6-8/group for **I-M.** * = p≤0.05, ** = p≤0.01, *** = p≤0.001. Error bars represent SEM.

Next, we investigated whether f-LDRT with *Sphk2* global KO could induce a greater anti-tumor response in the lymphoid and myeloid immune cells than *Sphk2* global KO alone. Indeed, f-LDRT in *Sphk2* global KO showed a significantly improved CD8^+^T cell to Treg ratio (**Figure 4H**) along with a significant increase in the fraction of CD8^+^T cells and a reduction in the fraction of Treg cells compared to *Sphk2* global KO (**Figures S3G,H).** Percentages of CD8^+^T cells, Tcon cells, and NK cells expressing IFNγ and Granzyme B (GrzB) significantly increased in the f-LDRT + *Sphk2* global KO arm compared to *Sphk2* global KO (**Figures 4I-K**). Percentage of CD4^+^FOXP3^-^ Tcon cells remained unchanged across all groups, while that of NK cells increased in the f-LDRT + *Sphk2* global KO arm over *Sphk2* global KO tumors (**Figures S3I-J**). Further, we found a significant shift towards the “M1-like” (anti-tumor) from the “M2-like” phenotype in TAMs and MGs in tumors (**Figure 4L,M**) and Mφs in cLNs (**Figure S3K**) after f-LDRT in *Sphk2* global KO compared to *Sphk2* global KO arm without f-LDRT. This response was accompanied by a significant increase in the fraction of NK cells in the cLNs secreting cytotoxic cytokines such as TNFα and GrB (**Figure S3L**) and overall NK cell percentage in the f-LDRT + *Sphk2* global KO arm compared to *Sphk2* global KO cLNs (**Figure S3M**). These findings support our hypothesis that f-LDRT has the potential to improve the immune susceptibility of G3MBs and further potentiate the anti-tumor activity of SPHK2- blockade.

### Combination treatment activates T and NK cells in G3MB tumors and cervical lymph nodes

We next tested whether the immune reprograming by f-LDRT with genetic perturbation of Sphk2, translated to survival benefit in mice. The combination of f-LDRT and Opaganib significantly improved median survival over each therapy alone in both MYC1 and MYC2 models (p<0.001) (**Figures 5A,B**). Median survival was 38 days (f- LDRT + Opaganib) compared to 14 days (f-LDRT) and 28 days (Opaganib) in MYC1 (**Figure 5A**). Median survival was 32 days (f-LDRT + Opaganib) compared to16 days (f- LDRT) and 21 days (Opaganib) in MYC2 (**Figure 5B**). To determine the effect of combination therapy on the tumor immune microenvironment, we profiled the lymphocytes from the time-matched MYC1 and MYC2 tumors and cLNs. The combination therapy enhanced the CD8^+^T cell to Treg cell ratio over control (**Figure 5C**) due to a significant increase in CD8^+^T cells and decrease in % Treg cells compared to control (**Figure S4A,B**) in both MYC1 and MYC2 tumors. There was a significant increase in the fractions of CD8^+^T, Tcon, and NK cells expressing IFNγ and GrB, with combination therapy compared to control (**Figure 5D-F**), although the percentage of Tcon cells and NK cells remained unchanged (**Figure S4C,D**). Overall CD45^+^ count significantly increased under combined therapy in MYC1 and MYC2 tumors compared to control (**Figure S4E**).

**Figure 5.**
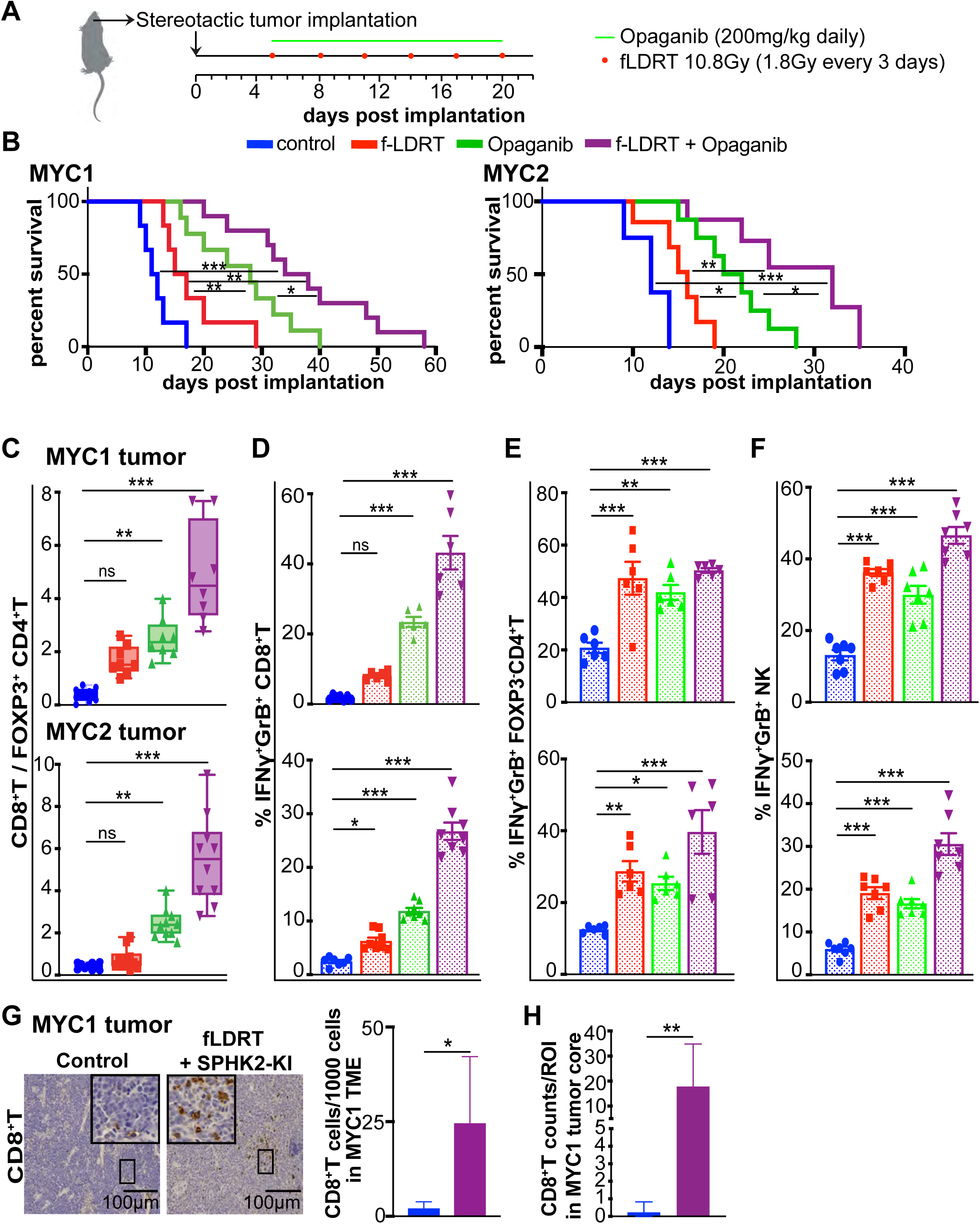
f-LDRT combined with pharmacological SPHK2 inhibition improved survival of mice bearing G3MB more than either therapy and reprogramed T and NK lymphocytes. **(A)** Schematic of f-LDRT and Opaganib treatment regimen. Mice with orthotopically implanted MYC1 or MYC2 tumor cells were randomized at day 5 after implantation (or when tumor volume is 3-4mm^3^). Mice were treated daily via oral gavage with 200mg/kg of Opaganib. Tumors in the cerebellum were irradiated with a 1.8Gy fraction every 3 days for 6 fractions. **(B)** Survival curves of mice bearing MYC1 and MYC2 tumors with indicated treatments. Median survival was compared across treatment groups. * = p≤0.05, ** = p≤0.01, *** = p≤0.001, n=15/group. **(C-F)** Flow cytometry of MYC1 tumors from mice with indicated treatments. Y axis denotes either ratio of populations **(C)** or percentage of population **(D-F)**. n=8-10/group for **C** and n=6-8/group for **D-F.** * = p≤0.05, ** = p≤0.01, *** = p≤0.001. Error bars represent SEM. **(G)** IHC staining and quantification for CD8+T cells in MYC1 tumors treated with Opaganib and fLDRT (n=2) relative to control (n=4). * = p≤0.05. Error bars represent SEM. **(H)** Quantification of CD8^+^T cell counts / ROI in the tumor core regions in MYC1 tumors treated with Opaganib and fLDRT (ROI n=10) relative to control (ROI n=13). Tumor core is defined as central tumor tissue, at least 100µm away from the normal cerebellum. ** = p≤0.01. Error bars represent SEM.

A critical limiting factor for immune response is the spatial distribution of effector cells in the tumor bed (Melero et al., 2014). We found that combined f-LDRT + Opaganib significantly improved CD8^+^, and CD3^+^ cells in the tumor core compared to control. (**Figures S4F-I**). CD8^+^ T cell count was significantly increased in f-LDRT + Opaganib - treated tumors compared to controls (**Figure 5G**). Randomly selected regions of interest (ROI) from IHC of tumor tissues showed significantly higher CD8^+^ T cell infiltration in the core of an f-LDRT + Opaganib-treated MYC1 tumor compared to the core of a control tumor (**Figure 5H**). To test if the T cells were clustered in certain intra-tumoral regions or distributed throughout the tumor area, we tracked CD8^+^ and CD3^+^ T cells from the edge of the tumor to the center. Interestingly, we noticed a consistent presence of T cells across the tumor area in the combined therapeutic arm, followed by the Opaganib arm, while the number of T cells steadily decreased in the control and f-LDRT groups with increasing distance from the tumor edge (**Figure S4J**). Finally, analysis of randomly chosen areas of interest on the tumor edge and centers showed increased CD8^+^ and CD3^+^ T cell numbers in both regions in the combined f-LDRT + Opaganib and Opaganib alone-treated groups compared to f-LDRT and controls, suggesting a tumor-wide recruitment of immune cells (**Figure S4K**). From our findings, we conclude that combining targeted therapy (SphK2 blockade) with modified standard of care (fractionated low-dose radiotherapy) can regress MB tumors cumulatively by activating T cells in the tumor microenvironment and facilitating T cell trafficking across the tumor area, including tumor bed, to generate a long-lasting anti-tumor response.

In concert with our tumor data, the ratio of CD8^+^T cells to Tregs in the cLNs was significantly improved with combination therapy compared to control (**Figure S5A**). This was due to a significant increase in the fraction of CD8^+^T cells in MYC1 and a decrease in the percentage of Tregs in both models (**Figure S5B,C**). The combined treatment also significantly increased the percentage of CD8^+^T cells secreting IFNγ and TNFα, as well as Tcon and NK cells secreting IFNγ and GrB in the cLNs compared to controls (**Figure S5D-F**). Tcon and NK cell percentages remained unchanged in the combination arm compared to control in both models (**Figure S5G, H**). CD45+ cells significantly increased in the combination arm only in MYC1 cLNs but remained unchanged in MYC2 cLNs (**Figure S5I**). Collectively, these data show that the added survival benefit is associated with increased T and NK activation in the tumor and cLNs.

### Combined treatment skews myeloid cells towards an anti-tumor phenotype in G3MB

Since myeloid-derived immune populations such as TAMs, Mφs, and DC-like cells play a significant role in anti-tumor immunity, we next asked whether combination therapy could reprogram those populations in G3MB tumors and cLNs. By immune profiling time-matched tumors via flow cytometry, we found that the TAMs and tumor-infiltrating brain tissue-resident MG populations were significantly skewed towards an M1-like versus M2- like phenotype in the combination treatment arm compared to control in both MYC1 and MYC2 tumors (**Figures 6A,B**). In addition to the commonly used cell surface markers, genes commonly associated with the M1-like phenotype (*Ccr6, Ccr9, Tnfα, Cxcr1, Cxcr3, Cxcl9* and *Inos*) and M2-like phenotype (*Tgfβ, Arg1, Ccl17, Ccl22, Vegf* and *Il10*) were respectively upregulated and downregulated in sorted TAMs from the combined therapeutic group over control in both tumor models (**Figure 6C**). The percentage of TAMs in all CD45+ cells did not change significantly in the combination group compared to control (**Figure S6A**). The MG percentage of all CD45+ cells only significantly increased in the combination arm of MYC1 compared to control (**Figure S6B**). The combined treatment group showed a significant increase in the fraction of MHCII+CD86+ DC-like cells compared to control (**Figure 6D**) without affecting the total DC-like cell percentage of all CD45+ cells (**Figure S6C**).

**Figure 6.**
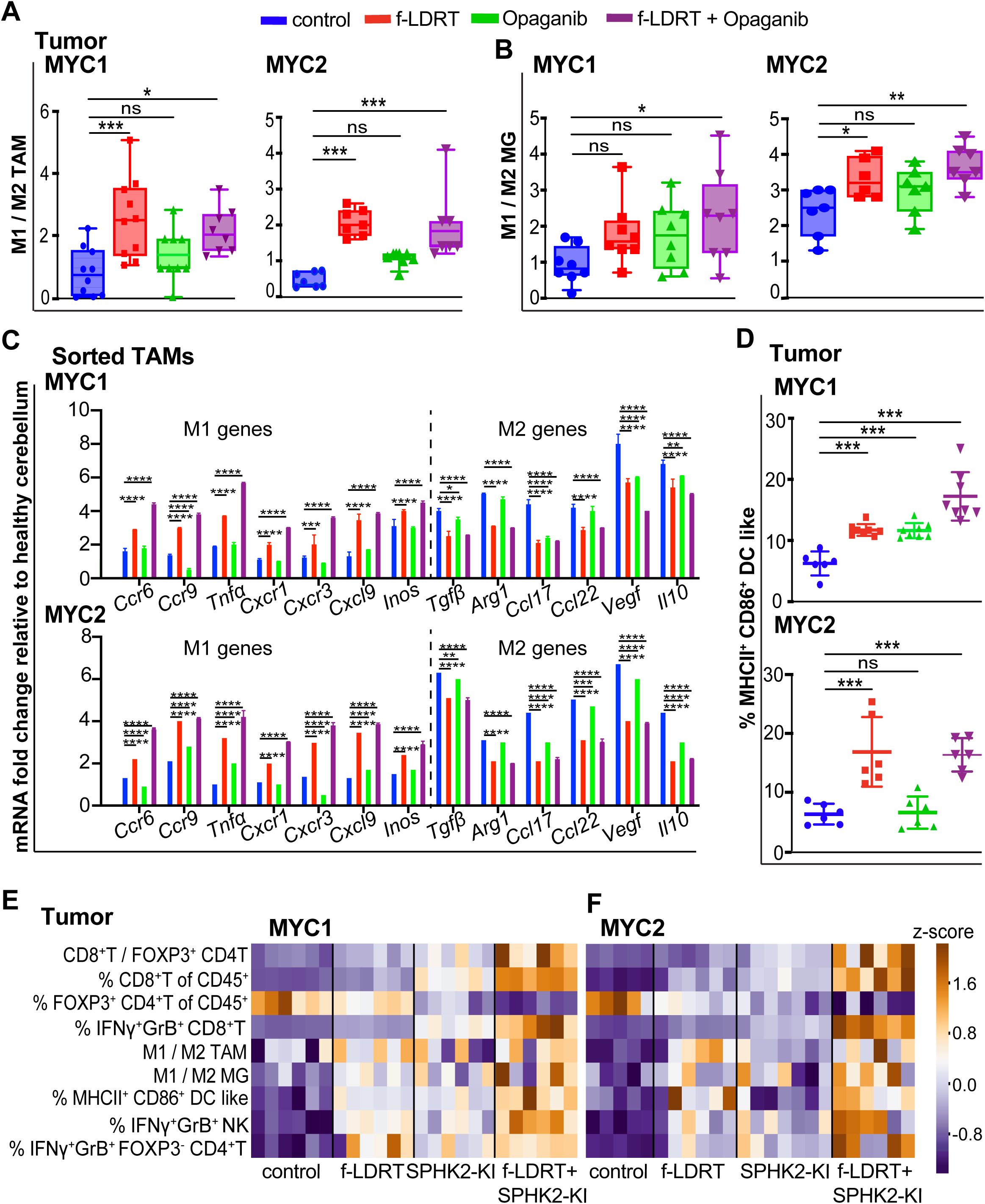
f-LDRT combined with SPHK2 inhibition is associated with myeloid cell repolarization in G3MB. **(A,B)** Flow cytometry analysis of MYC1 and MYC2 tumors from mice treated with indicated regimens. M1 / M2 ratio is shown for TAMs **(A)** and MG **(B)**. n=8-10/group for **A** and n=6-8/group for **B**. * = p≤0.05, *** = p≤0.001, **(C)** Expression of indicated genes using qPCR in FACS-sorted TAMs (CD45highGr-1^-^ CD11c^-^CD11b^+^F4/80^+^) from MYC1 and MYC2 tumors. Tumors were collected from mice with indicated treatments. Expression in healthy cerebellum was used for normalization. n=3/group. *** = p≤0.001, **** = p≤0.0001. Error bars represent SEM. **(D)** Flow cytometry analysis of MYC1 and MYC2 tumors from mice treated with indicated regimens. Percentage of MHCII^+^CD86^+^ DC-like cells (CD45^+^CD11c^+^CD11b^-^Gr1^-^F4/80^-^) are shown. n=6-8/group. *** = p≤0.001. **(E, F)** Heatmap showing flow cytometry analysis of MYC1 **(E)** and MYC2 **(F)** tumors for indicated populations Tumors were collected from mice with indicated treatments.

In line with the tumor data, Mφs in the cLNs were skewed towards an M1 phenotype in the combination arm compared to control (**Figure S6D**). Mφs as a percentage of CD45^+^ cells did not vary under dual treatment compared to control (**Figure S6E**). The percentage of APC DC-like cells was significantly increased in the combination arm over control (**Figure S6F**).

To visualize the complexity of the immune microenvironment of G3MB and the subsequent reprogramming under combination versus monotherapies, we generated a heat map summarizing the flow cytometry data as z-scores of different immune populations, or their ratios (**Figure 6E,F**). Collectively, these data show that, while each monotherapy can reprogram key players of the tumor and cLN immune microenvironment, combining treatments lead to an additive anti-tumor response in G3MB.

### Combined treatment stimulates systemic anti-tumor immunity in mice with G3MB

Since tumors are known to affect both local and systemic immunity, we asked whether combined treatment extended anti-tumor immunity to the peripheral sites such as the spleen and peripheral blood, and led to anti-tumor memory after 15 days of treatment. Opaganib with or without f-LDRT showed a significantly higher ratio of CD8^+^T cells to Tregs in the spleen (**Figure 7A**). This was associated with a significant increase in CD8^+^T cell percentage in all groups compared to control (**Figure S7A**). Treg percentage or CD45^+^ immune cell counts were not affected in any arm over control (**Figure S7B,C**). Opaganib with or without f-LDRT significantly increased the percentage of CD8^+^T cells secreting IFNγ and TNFα in the spleen compared to control (**Figure 7B**). f-LDRT with or without Opaganib significantly increased the percentage of NK cells secreting IFNγ and TNFα in the spleen compared to control (**Figure 7C**) while NK cell fraction remained unchanged (**Figure S7D**). Interestingly, the percentage of CD44high CD62L^−^ effector memory CD8^+^T cells increased most significantly after combined therapy (**Figure 7D).**

**Figure 7.**
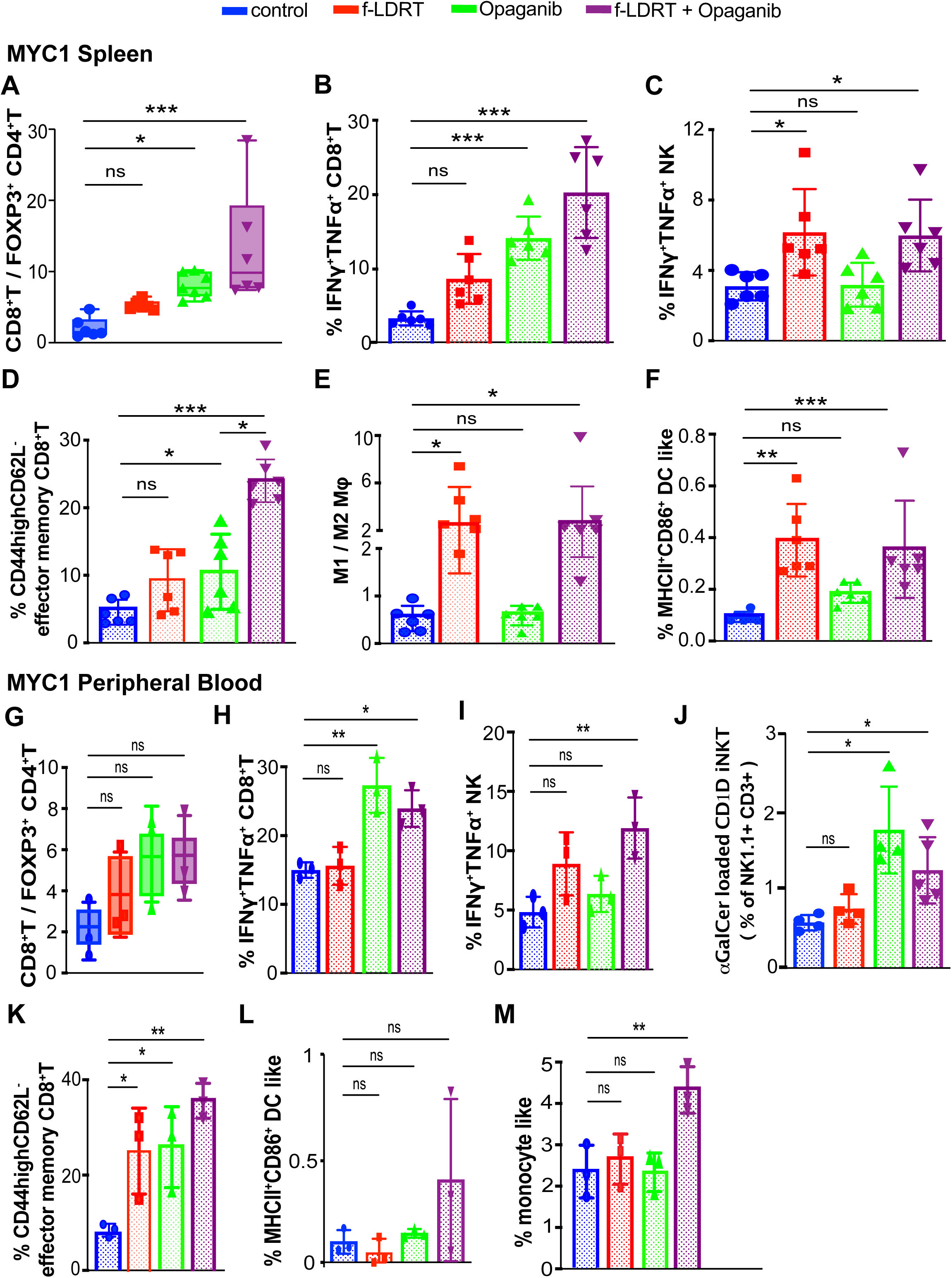
f-LDRT combined with Opaganib stimulates systemic anti-tumor immunity in mice with G3MB. **(A-F)** Flow cytometry of MYC1 spleen tissues from mice with indicated treatments. Y axis denotes either ratio of populations **(A, E)** or percentage of population **(B-D, F)**. n=5- 7/group for **A**, n=5-6/group for **B**, n=6/group for **C-F**. * = p≤0.05, ** = p≤0.01, *** = p≤0.001. Error bars represent SEM. **(G-I)** Flow cytometry analysis using peripheral blood from mice bearing MYC1 tumors and treated with indicated regimens. Y axis denotes either ratio of populations **(G)** or percentage of population **(H, I)**. n=3/group for **G-I**. * = p≤0.05, ** = p≤0.01, *** = p≤0.001. Error bars represent SEM. **(J)** Flow cytometry analysis using peripheral blood from mice bearing MYC1 tumors treated with indicated regimens. Percentages of iNKT (NK1.1^+^CD3^+^) cells are shown on the Y axis. n=4/group. * = p≤0.05, *** = p≤0.001. Error bars represent SEM. * = p≤0.05, *** = p≤0.001. Error bars represent SEM. **(K-M)** Flow cytometry analysis using peripheral blood from mice bearing MYC1 tumors and treated with indicated regimens. Y axis denotes percentage of indicated populations. n=3/group for **K-M**. * = p≤0.05, ** = p≤0.01. Error bars represent SEM.

In line with the tumor data, f-LDRT with or without Opaganib significantly skewed Mφs towards an M1-like phenotype and increased the percentage of APC DC-like cells compared to controls (**Figure 7E,F**). While the percentage of Mφs remained unchanged (**Figure S7E**), the percentage of DC-like cells only significantly increased in f-LDRT arm compared to control (**Figure S7F**). In peripheral blood (PB), the CD8^+^T cells to Tregs ratio did not significantly change in any arm (**Figure 7G**). This was due to a lack of significant change in the CD8^+^T cell and Treg percentages across all groups compared to control (**Figure S7G,H**). CD45+ immune cell count was significantly reduced only in f-LDRT (**Figure S7I**). Opaganib with or without f-LDRT significantly increased the percentages of CD8^+^T cells secreting IFNγ and TNFα compared to control (**Figure 7H**). Only the combination therapy significantly increased the percentage of NK cells secreting IFNγ and TNFα compared to control (**Figure 7I**). The NK cell percentage was only significantly increased under f-LDRT (**Figure S7J**). Presentation of α-Galactosyl Ceramide (α-GalCer) was associated with increased frequencies of CD1D-restricted iNKT cells in the Opaganib and combination groups, whereas f-LDRT alone had little effect, suggesting a predominantly drug-driven response. **(Figure 7J).** The memory CD8^+^ T cell fraction increased significantly in all arms compared to control (**Figure 7K**). The percentages of APC DC-like cells and overall DC-like cells did not significantly change in any arm compared to control (**Figures 7K and S7K**). The percentage of monocytes significantly increased only in the dual therapy arm compared to control (**Figure 7M**). These findings provide evidence that the combination of f-LDRT and Opaganib as well as monotherapies, can induce systemic immune stimulation in mice with G3MB.

## Discussion

Recent studies indicate that tumor-subgroup-specific genes and immune microenvironmental signatures could be leveraged to develop targeted molecular therapies for G3MB (Menyhárt et al., 2019; Park et al., 2019; Pham et al., 2016). Despite these advances, there is no validated targeted therapy for G3MB.

Our study reveals for the first time that SPHK2 is a molecular driver of tumorigenesis in G3MB and serves as an upstream regulator of cMYC, pERK and pSTAT3. SPHK2 has been linked to apoptosis and cell death, with reports proposing different potential modes of how it can drive oncogenesis (Song et al., 2019; Xu et al., 2018). Genetic silencing of *Sphk2 in* tumors increased expression of *e2f*, a known tumor suppressor and activator of *p53*, (Li et al., 2018; Polager & Ginsberg, 2009). *Sphk2* knockdown tumors also had reduced expression of *Fgfr3*, which was recently shown to promote MB proliferation (Holzhauser et al., 2020). Inhibiting SPHK2 reduced the cMYC level, a master transcriptional regulator of G3MB and an inducer of immunosuppression. This is in line with two reports showing that Opaganib inhibited cMYC in acute lymphocytic leukemia and multiple myeloma (Venkata et al., 2014; Wallington-Beddoe et al., 2014). Thus, our findings support SPHK2 inhibition as a viable strategy to inhibit oncogenic cMYC signaling, frequently amplified in G3MB patients.

We show that Opaganib, an SPHK2 inhibitor, is selectively cytotoxic to MYC1 G3MB cells, which have elevated levels of *Sphk2,* compared to the cerebellum. Opaganib is well-tolerated in tumor-bearing mice with no significant weight loss. This finding is in line with other studies that found Opaganib to be safe and non-toxic in mice and well-tolerated in adult patients at a dose of 500mg daily for 28 days (Britten et al., 2017; French et al., 2010). This could be attributed to the fact that Opaganib competes with sphingosine to bind to SPHK2, preventing the conversion of sphingosine into S1P. This property strongly reduces the off-target effects of Opaganib on other protein kinases (Britten et al., 2017; French et al., 2010). Thus, targeting SPHK2 with Opaganib can be particularly beneficial for treating pediatric patients with tumors expressing high SPHK2 while reducing off-target toxicity.

MBs are known to be immunosuppressive tumors due to the lack of lymphocyte infiltration, MHCI expression, and acquired expression of granzyme inhibitors (Spranger, 2016; Vermeulen et al., 2018). Yet, key molecular drivers of immunosuppression in G3MB have not been identified yet. Comprehensive information about the spatial distribution of immune cells during MB progression is also lacking. SPHK2 has been shown to promote glioma growth by skewing TAMs towards an M2-like (pro-tumor) phenotype (J. Liu et al., 2018). However, the specific role of SPHK2 in suppressing anti-tumor immunity in G3MB is not known. Our study shows that targeting SPHK2 alleviates immunosuppression in pediatric G3MB TME via multiple mechanisms. SPHK2 WT tumors have poor CD8^+^T cells infiltration and activity along with increased Tregs. Genetic and pharmacological targeting of sphingosine kinase 2 significantly improved the ratio of CTLs to Tregs in the TME. *Sphk2* global KO cLNs had reduced expression of *Tgfb2*, a known immunosuppressor and tumor promoter in MB (Gate et al., 2014). Blocking SPHK2 increased the percentage of CD8^+^ T, NK, and NKT cells secreting inflammatory cytokines. The fact that cytotoxic T cells were activated under SPHK2 inhibition despite the proven lack of expression of MHCI on the tumor cells could be attributed to the alternate activation of NKT cells via CD1d. In mice, intracranial injection of NKT cells can result in the regression of CD1D+MB in mice (D. Liu et al., 2013). Blocking SPHK2 failed to significantly improve the ratio of M1 (anti-tumor) to M2 (pro-tumor) in TAMs and MGs in the G3MB TME (percentages of MHCII^+^CD86^+^ APCs in the TME and cLNs), which was consistent with findings that G3MBs evade immune surveillance via downregulating immune priming by APCs (Garancher et al., 2020; Haworth et al., 2014; Pham et al., 2016; Vermeulen et al., 2018). These data indicate impaired immune priming, indicating the need for an added strategy to harness the benefit of myeloid cell reprogramming.

Emerging evidence indicates that radiation doses and fractions can differentially affect the TME (Demaria & Formenti, 2012; Gameiro et al., 2014). Radiation can mediate cell stress, which can trigger DAMP-driven proinflammatory micro-environment required for an efficient innate immune response (Constanzo et al., 2021). In the clinic, craniospinal irradiation (CSI) is usually given via dose fractions over ∼13 to 20 days. Dose per radiation fraction and total treatment time impact the clinical response to radiotherapy (Yock et al., 2016). We found that a radiation dose of 10.8Gy given as a fractionated dose of 1.8Gy every 3 days to the tumor-bearing area of the cerebellum induced the upregulation of *Cd1d,* with a gradual increase over 6 fractions spaced 3 days apart. f- LDRT caused cytosolic genomic DNA release, which could have happened both from the context of malignant and immune cells in the TME activating the cGAS–STING signaling cascade in cDC1, which are well-known cross-presenting APCs (Embgenbroich & Burgdorf, 2018). This was in line with increased MHCII expression on multiple other APCs such as cDC2 and TAMs, under f-LDRT but not in the Sphk2 global KO, indicating that this is an f-LDRT-driven effect in the tumors. However, f-LDRT alone was incapable of controlling tumor progression despite the improved potential of APCs to present tumor-specific antigens. The latter is in line with clinical results showing that standard radiotherapy alone is not sufficient to overcome immunosuppression. We show here how f-LDRT itself and in combination with Opaganib can be exploited for their immuno- stimulatory benefit to reprogram the immune landscape in the TME while curtailing radiation-associated lymphopenia, severe weight loss, skin rashes, and hair loss. The combined therapy can reprogram diverse lymphoid and myeloid immune landscapes in the TME to enhance the anti-tumor response and survival beyond either monotherapy. Additionally, we found an increased influx of CD8^+^T cells into the tumor core under combined therapy, suggesting a tumor-wide immune response.

Systemic tissues, such as peripheral blood and spleen, are frequently used as surrogate sources to assess immune reprogramming patterns occurring within the tumor microenvironment and other peripheral organs. (Larsson et al., 2019; Spitzer et al., 2017). We found that immune cells in the spleen of G3MB mice are immunosuppressed. The combination, as well as monotherapies, could alleviate immunosuppression in the spleen and induce inflammatory cytokines in CD8+T and NK cells in the spleen and peripheral blood. Interestingly, although combination therapy significantly improved markers of effector memory response in CD8+T cells, eventually all mice succumbed to the tumors even after a prolonged survival benefit. Hence, future investigations need to address the acquired resistance.

Our study has several limitations. The molecular classification of the patient tissues from the commercially available patient microarray was not available. Therefore, LCA histology was chosen for IHC staining of SPHK2 because of its frequent association with G3MB. The effects of the proposed combination therapy may extend to other immune cell populations, such as myeloid-derived suppressor cells (MDSCs). MDSCs, long known for their immune-suppressive properties, are present in the TME of murine G3MB (Pham et al., 2016). Fractionated radiation of 15Gy has been shown to induce monocytic (M) and polymorphonuclear (PMN) MDSC infiltration into prostate tumors (Wu et al., 2013). Therefore, further elucidation of the phenotypes of MDSCs might help develop a better understanding of their role in inducing resistance to this combined treatment modality in G3MB. In our study, we also did not examine the potential role of blocking S1PR1. Surface expression of the S1PR1 receptor facilitates T-cell egress from the spleen, LNs, and thymus into the circulation. A S1PR-agonist that causes downregulation of surface S1PRs, FTY720, is shown to confine T-cells in the LNs and prevent their circulation (Aoki et al., 2016). Our findings of significantly increased CD8^+^T cell percentage in the tumor and unchanged CD8+T cell percentage in peripheral blood, as well as unaltered level of S1PR1 protein under Opaganib, indicate a different mechanism of action for SPHK2 inhibition compared to FTY720.

In summary, we show that SPHK2 in cancer cells can drive G3MB initiation, tumor progression and immune evasion, creating a tumor-supportive microenvironment. Blocking SPHK2 with a low MW, brain-permeable inhibitor induces a cascade of anti-proliferative, apoptotic, and immune-stimulatory signaling, making it an attractive therapeutic target. Our proposed combinatorial therapeutic approach reprograms the immunosuppressive TME into an immunostimulatory milieu. We show that the immuno- stimulatory benefit of radiation can be enhanced by modifying the total dose, dose fractions, and the durations between fractions without inducing radiation-associated lymphopenia. Importantly, administration of Opaganib with fractionated low-dose radiotherapy (10.8Gy) alleviates immunosuppression additively and regresses tumors beyond individual monotherapies without significant lymphopenia or weight loss in immunocompetent mouse models of G3MB. Our findings provide a compelling rationale and supporting data to test this combination in patients with G3MB.

## Materials and methods

### KEY RESOURCES TABLE

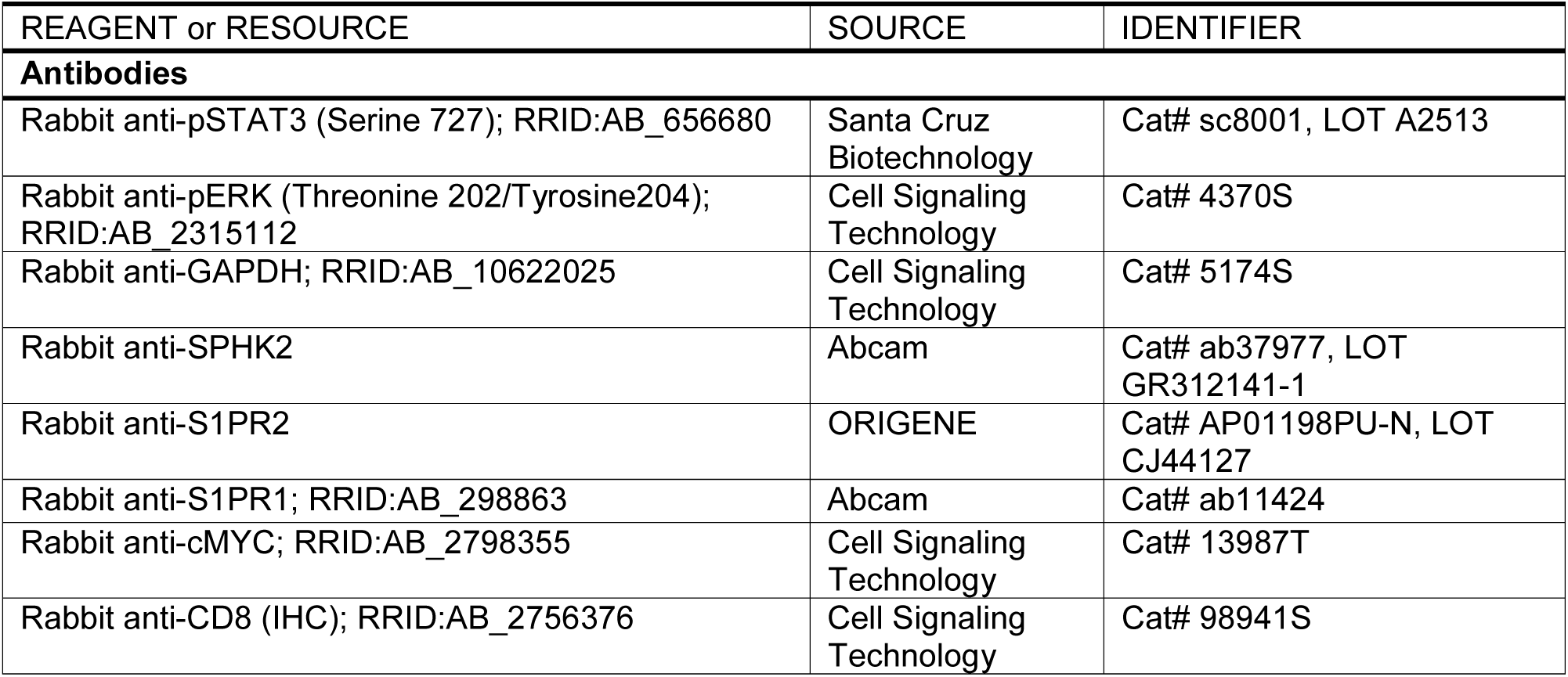
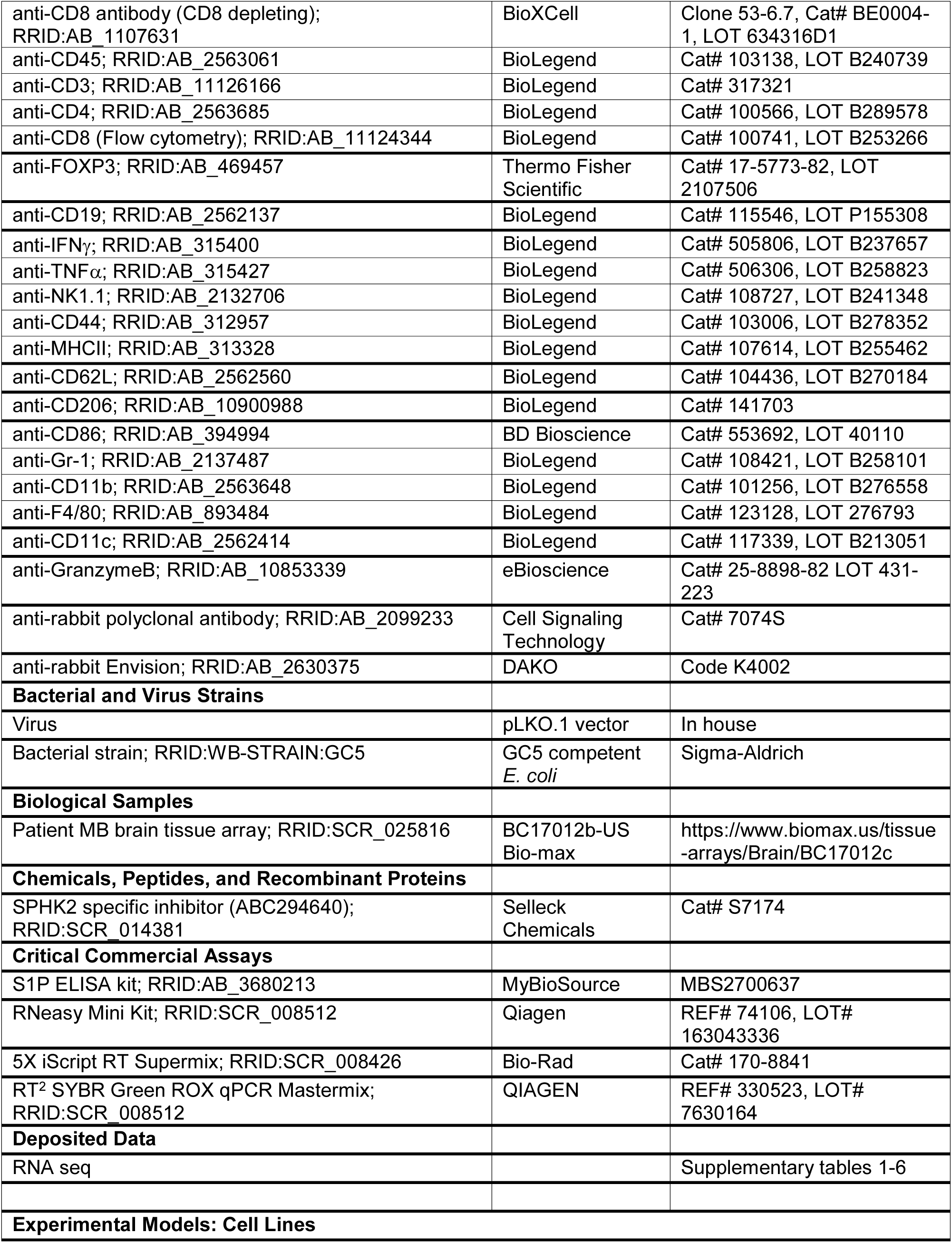
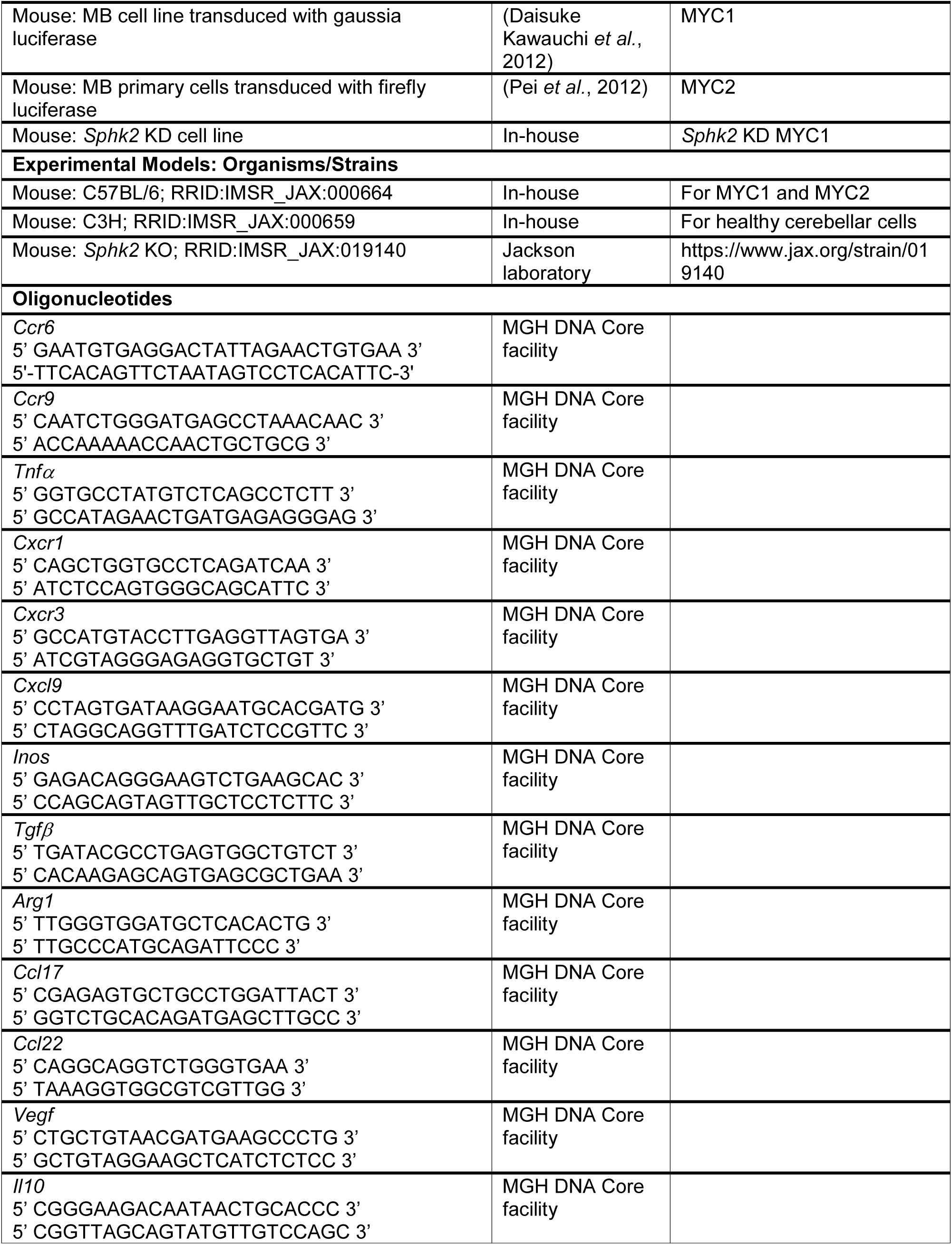
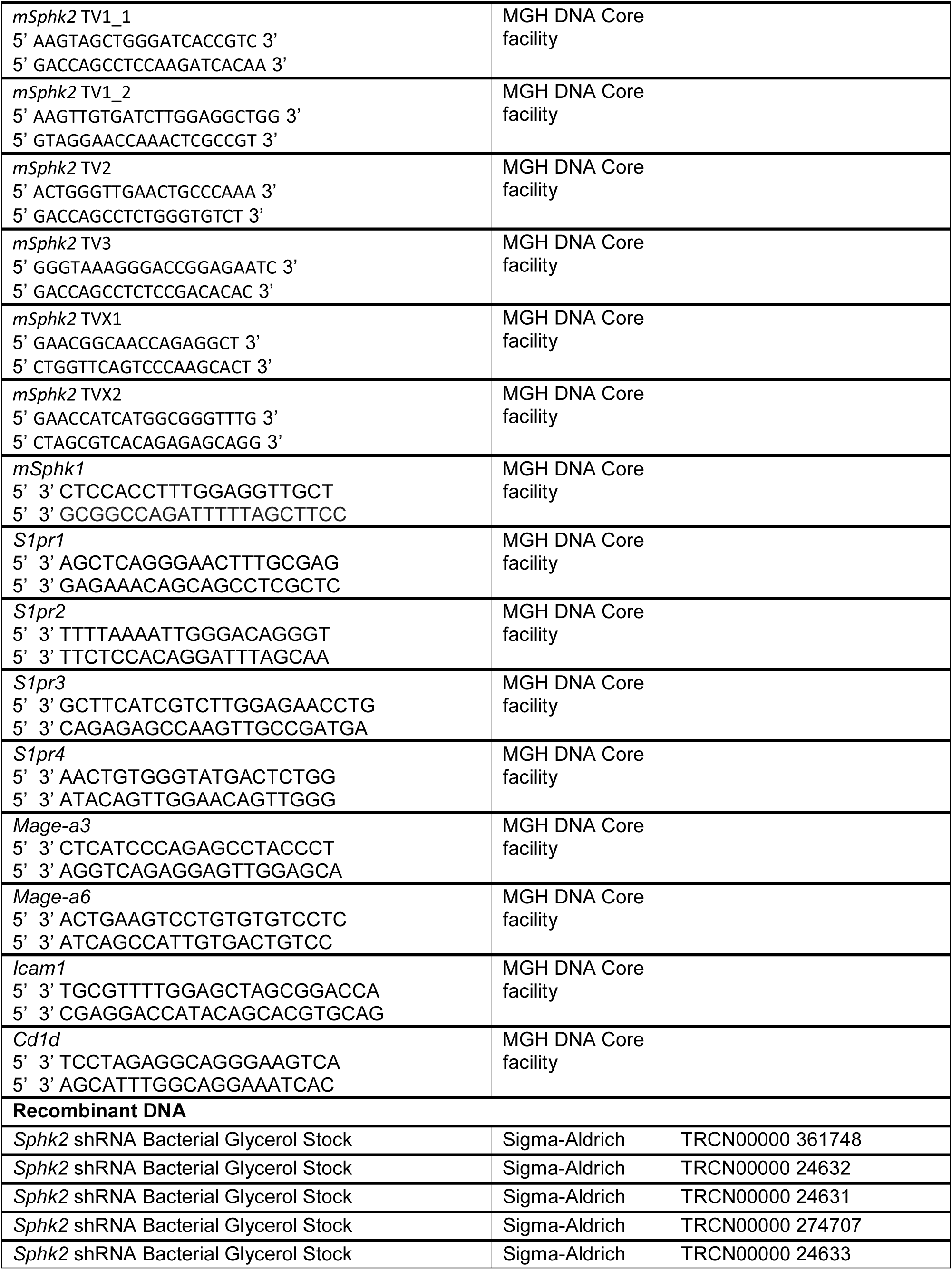
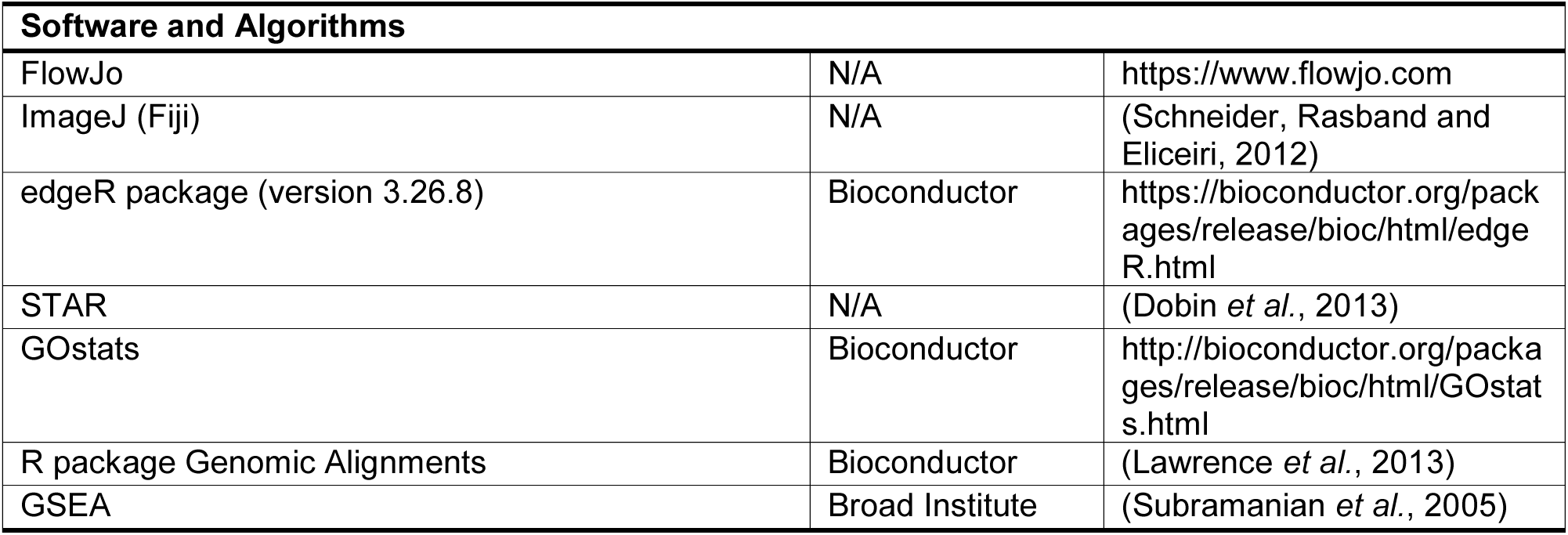

### Mice

*Sphk2* KO mice were purchased from Jackson Laboratories (https://www.jax.org/strain/019140). C57BL/6 and C3H mice were bred in-house.

### Primary tumors, cell lines and primary cell culture

We obtained two murine MB primary tumors recapitulating the MYC-driven subtype, MYC1 (D Kawauchi *et al*., 2012) and MYC2 (Pei *et al*., 2013). MYC1 tumor cells were received as frozen vials as a gift from Martine F Roussel’s lab, St. Jude Children Research Hospital (Daisuke Kawauchi *et al*., 2012) and was engineered to secrete Gaussia luciferase (g-luc) in our lab as done before (Chung *et al*., 2009). MYC2 mouse primary tumor cells (wild type) engineered to secrete firefly luciferase were received as frozen vials as a gift from Dr. Robert Wechsler-Reya’s lab, Sanford Burnham Prebys Medical Discovery Institute, La Jolla, CA (Pei *et al*., 2013). Only MYC1 was grown *in vitro* while both tumors were propagated in C57BL/6 mice up to F2 generation.

*Sphk2* was knocked down in MYC1 cell line (*Sphk2* KD) in-house (**refer to knockdown of *Sphk2* in MYC1 cell line section**). Primary culture of healthy cerebellar cells was established by isolating C3H mice brain tissue. Tissues were disintegrated and cells were cultured and passaged once before used for experiments. MYC1, *Sphk2* KD MYC1 and healthy cerebellar cells were maintained in Neurobasal medium (Life Technologies) with (complete) or without (starving) factors – 1X B27 (Life Technologies), 1X N2 (Life Technologies), 1% v/v L-glutamine (Life Technologies), 25mg/ml EGF (Peprotech), 25mg/ml FGF (Peprotech) and 0.78% BSA (Sigma). 293T cells were used for viral production during knockdown of *Sphk2* and were maintained in DMEM media (Gibco) with 10% fetal bovine serum (Sigma).

### Transduction of g-luc in murine cell lines

MYC1 cell line was transduced with Gaussia luciferase (lentiviral CSCW-g-luc; MGH DNA Vector Core) (Chung *et al*., 2009).

### Knockdown of *Sphk2* in MYC1 cell line

Five ampicillin-resistant bacterial stocks carrying plasmids coding for *Sphk2* shRNA with distinct sequences were ordered from Sigma-Aldrich as follows TRCN00000 361748, TRCN00000 24633, TRCN00000 24632, TRCN00000 274707, TRCN00000 24631. 293T cells were transfected with a construct that included a mixture of packaging vector, transfer (plasmid coding for shRNA), FuGENE and optimen. Scrambled shRNA was used as a control. MYC1 cells in culture were transduced with lentiviral particles and 22 µl of polybrene. Puromycin-selected MYC1 cells were cultured for several passages. RNA was isolated using the RNeasy Mini Kit (QIAGEN) from the transduced cells, and cDNA was synthesized with the Bio-Rad 5X iScript RT Supermix (Cat# 170-8841) in accordance with the manufacturer’s instructions. RT-PCR using SYBR green master mix (SYBR® Green Supermix Bio-Rad) was performed to determine the level of *Sphk2* in different transcript variants. The complete list of primers designed in-house using NCBI BLAST and obtained from MGH DNA Core facility is in the **Supplementary Methods section.** Fold changes in mRNA expression were calculated using the 2-ΔΔCt method by the software (MxPro-Mx3000P) and normalization was done using the housekeeping gene GAPDH. Sphk2 knockdown cells from shRNAs TRCN00000 361748 (shRNA1), and TRCN00000 24632 (shRNA2) with 60-80% knockdown efficiency were selected. TRCN00000 24631 (shRNA3) was discarded due lowest knockdown efficiency. *Sphk2* knockdown cells from shRNAs TRCN00000 24633 and TRCN00000 274707 with above 80% knockdown efficiency were not viable since they failed to develop neurospheres. *Sphk2* KD MYC1 tumor cells (mix of shRNA1 and shRNA2) were expanded and used for cerebellar implantation.

### Clinical samples and histopathology arrays and scoring

Patient MB brain tumor tissues and blood plasma samples (MB and healthy) were obtained in accordance with the Institutional Review Board (IRB) at the Massachusetts General Hospital under protocol number 05-300. Patient arrays (BC17012b-US Bio-max) were bought from US Biomax, Inc (https://www.biomax.us/tissue-arrays/Brain/BC17012c).

### Immunohistochemistry

Tumor tissues were harvested from MYC1 and MYC2 mice post-implantation. IHC was performed with 5µm formalin-fixed (24 hours) paraffin-embedded sections (patient – MB brain; immunocompetent models - MYC1 and MYC2). Slides were de-paraffinized, retrieved with specific antigen retrieval solutions, blocked with 5% H2O2 for 10 minutes and 5% normal donkey serum (NDS) for 1 hour followed by overnight incubation with respective primary antibodies and 30 minutes incubation with anti-rabbit Envision (DAKO) secondary antibody. Antigen retrieval solutions, primary antibody dilutions and diluent - 0.05% trypsin (Sigma) for anti-SPHK2 (1:30, 5% NDS), citrate pH9 for pSTAT3 ser727 (1:100, 5% NDS) and 1mM EDTA (Promega) pH8 for CD8 (1:50, 5% NDS). Washing was performed 3 times for 3 minutes between each step using 1X phosphate buffer saline. Slides were scanned using ZEISS Axioscanner. Images were analyzed using ImageJ – Fiji.

### Immunohistochemistry analysis

CD8^+^ stained tumor regions were manually segmented using ImageJ. Three kinds of analysis were performed: immune cell infiltration (CD45+, CD3+ and CD8+), core versus periphery (CD3+ and CD8+) analysis, and trafficking distance analysis (CD3+ and CD8+). The positive and total cell (based on hematoxylin nuclei stain) counts were identified by adjusting the HSB color threshold and analyzing particles. CD8^+^ fraction was calculated as number of CD8^+^ counts per 1000 cells. Region at least 100µm away from tumor edge adjacent to healthy area was considered as tumor core. Random regions of interest (ROI) in the tumor core were manually segmented and CD3+ or CD8^+^ cells were calculated as counts per ROI in tumor core. For trafficking distance analysis tumor regions were segmented and the boundaries between healthy tissue and tumors were manually demarcated. Positively stained cells were identified within the tumor region, and the invasion distance was calculated as the shortest distance between the positive cell and the tumor-healthy boundary. A final histogram was generated showing the number of positively stained cells per µm from the periphery. This entire analysis was conducted using custom code written in MATLAB (Mathworks).

SPHK2 expression in patient medulloblastoma brain tissue was scored using stained cell percentage in the nuclei of tumor cells by in-house pathologists (<20%=0, 20-50%=1, 50-80%=2 and >80%=3).

### qPCR

qPCR was performed to investigate the fold change in expression of *Sphk1, Sphk2, S1pr1, S1pr2, S1pr3, S1pr4, Mage-a3*, *Mage-a6*, *Icam1* and *Cd1d* in MYC1 tumors and *Ccr6, Ccr9, Tnfα, Cxcr1, Cxcr3, Cxcl9, Inos, Tgfβ, Arg1, Ccl17, Ccl22, Vegf* and *Il10* in flow-sorted TAM population in MYC1 and MYC2 tumors. TAMs were sorted by BD FACSAria™ Fusion (refer to FACS section). Tissues were harvested and homogenized with Kimble™ Kontes™ Pellet Pestle™ Cordless Motor homogenizer (DWK Life Sciences 7495400000) in RLT buffer and run through QIAshredder spin columns (QIAGEN). RNA was extracted using RNeasy Mini Kit (Qiagen) with DNase I treatment and cDNA was prepared using 5X iScript RT Supermix (Bio-Rad). SYBR green master mix (SYBR® Green Supermix Bio-Rad) was used for qPCR. All kits were used as per manufacturer’s protocol.

### ELISA

ELISA was performed to detect S1P levels in circulating blood plasma (patient – MB and healthy, murine – MYC1). Peripheral blood of MYC1 mice was collected using cardiac puncture and serum extracted using PBMC fractionation. ELISA was carried out using an S1P ELISA kit (MyBioSource) as per the manufacturer’s instructions and was measured using iMark microplate reader (BioRad).

### Tumor implantation and measurement

Approximately 25,000 MB tumor cells (MYC1, MYC2 and *Sphk2* KD MYC1) were re-suspended in Bambanker solution (Thermo fisher Scientific) and stereotactically implanted in mice (wild type C57BL/6 and *Sphk2* global KO C57BL/6) as described in (Snuderl *et al*., 2013). Mice were subcutaneously injected with 0.1mg/kg body weight of Buprenorphine (Buprenex, Patterson Veterinary) 15 minutes prior surgery, anesthetized with a mixture of 90 and 9 mg/kg body weight of Ketamine (Ketaset, Patterson Veterinary) and Xylazine (AnaSed, Patterson Veterinary) respectively and injected with 0.1 mg/kg body weight of Buprenorphine the next day. All mice were 4-5 weeks in age at the time of implantation.

MYC1 and *Sphk2* KD MYC1 tumor volume was correlated simultaneously via ultrasound and secreted g-luc levels in tail vein blood, measured via luminometer with coelenterazine (CTZ) as the substrate (Askoxylakis *et al*., 2017). MYC2 tumor volume was measured by whole body imaging. Briefly, mice were injected intraperitoneally with 150ul of D-luciferin (LUCK-2G Gold Bio- Products) and anesthetized after 10 minutes with ketamine. Whole body imaging was performed using ICIS spectrum whole body imager and estimated by correlation of firefly luciferase mean fluorescence (counts/s/cm2) with tumor size (mm3) (Pei et al., 2012).

### Treatment regimen for mice

Drug - Tumor bearing mice were treated with 200mg/kg body weight of SPHK2-SI, ABC294640 via oral gavage daily post 5 days of implantation.

Irradiation – Mice were irradiated with 1.8Gy of local X-radiation using RAD 320 irradiator (Precision X-RAY INC. N.Branford, CT) at the site of tumor implantation (3.76 Gy per min; 10 × 10 mm field size).

### Mass Spectrometry imaging (MSI) of glycolipids in MYC1 and MYC2 tumors

MALDI mass spectrometry was performed to detect glycolipids in MYC1 and MYC2 tumors at day 8. Tissue sections (12 μm) were cut from brain biopsy specimens using a Leica CM1850 cryostat (Walldorf, Germany) and thaw-mounted onto standard glass microscope slides (as Blanc et al, Anal Chem, 2018).

For the MALDI-MSI analysis, diaminonaphthalene (DAN, Sigma-Aldrich, St. Louis, MO) matrix, prepared at 5 mg/mL in acetone/water (7:3) was applied to the tissues via a TM-Sprayer automated MALDI tissue-prep device (HTX Technologies, Chapel Hill, NC) under the following optimized conditions: a 0.05 mL/min flow rate, a 60 °C nozzle temperature, and a 1.3 mm/s raster speed with 20 passes over the tissue.

MALDI-MSI acquisition was performed using a MALDI LTQ Orbitrap XL mass spectrometer (Thermo Fisher Scientific, Bremen, Germany) with a resolution of 60 000 at m/z 400 (full width at half maximum). The imaging data were acquired in full-scan mode to maximize sensitivity. Spectra were acquired in negative mode, across the mass range of m/z 200–2000 with a laser energy of 20 μJ and five shots per position (one microscan per position).

2D ion images were generated using Thermo ImageQuest software (v1.01). Normalized ion images of lipid signal were generated by dividing with the Total Ion Count (TIC) and abundance maps of the following were created: red: Ganglioside GM2 (d34:1) – m/z 1493.946 which colocalizes with tumor; green: Sulfated Hexose Ceramide (d42:2) – m/z 888.624 which colocalizes with white matter; blue: Docosahexaenate (DHA) - m/z 327.233, fatty acid which colocalizes with gray matter.

### Mass Spectrometry of ABC294640 in MYC1 and MYC2 tumors

Mass spectrometry of MYC1 and MYC2 brain tissues was performed at to determine the concentration of ABC294640. All reagents and materials coming into contact with tissues were pre-chilled on dry ice for at least 30 minutes. Tissues were weighed out into pre-chilled Eppendorf Snap-Cap microcentrifuge safe-lock tubes (Fisher Scientific) followed by the addition of 600 ul of 60% methanol (Alfa Aesar) and 400 ul of HPLC-grade chloroform. After vigorous vortexing, two 5 mm stainless steel beads (Qiagen) were added to each tube and loaded on the TissueLyzer at 50 Hz for 3 minutes. Beads were then removed, and samples were centrifuged at 13,000 rpm for 15 minutes. The aqueous layers were transferred to fresh Eppendorf tubes and evaporated in a Savant™ SPD131DDA SpeedVac™ (Thermo Fisher) at 4°C overnight.

The detection of ABC294640 on an Agilent 6470 series triple quadrupole mass spectrometer was optimized using the built-in MassHunter Optimizer program, revealing a precursor ion of m/z 379.16 and four fragment ions (in decreasing abundance: m/z = 78.1, 42.2, 133 and 105.1). ABC294640 was resolved using reverse phase ion-pairing chromatography on an Agilent 1290 Infinity II Series LC and detected in a full scan (negative mode) by the aforementioned mass spectrometer. The chromatography method was adapted from a targeted dMRM database and method from Agilent Technologies. Dried samples were resuspended in Buffer A (97% deionized water, 3% methanol, 10 mM tributylamine, 15 mM glacial acetic acid, pH 5.5) and injected over a Buffer A-equilibrated ZORBAX Extend-C18 column (2.1 x 150 mm, 1.8 µM). Samples were eluted for 2.5 minutes with 0% Buffer B (10 mM Tributylamine, 15 mM Glacial Acetic Acid in 100% Methanol), followed by a linear gradient of 0-20% Buffer B (5 minutes), 20-45% (5.5 minutes), 45- 99% (7 minutes) and 99% for 4 minutes. Samples were ionized in negative polarity using an Agilent Jet Stream Source with the following MS source parameters: nebulizer = 45 psi, capillary voltage = -2000V, nozzle voltage = 500 V, sheath gas temperature = 325°C, sheath gas flow = 12 L/minute, gas flow = 13 L/minute, and gas temperature = 150°C.

### Viability assay

MYC1 tumor cells and healthy cerebellar cells were seeded, incubated in starving media for 24 hours and treated with 0.1µM, 0.5µM, 0.7µM and 1µM of SPHK2-SI, ABC294640 (Selleck Chemicals, Cat# S7174), stocks were prepared in DMSO. MTT assay was performed to determine the viability and was measured using iMark microplate reader (BioRad).

### Western Blot

MYC1 cell line was seeded, incubated in starving media for 24 hours and treated with 0.1µM, 0.5µM, 0.7µM and 1µM of SPHK2-SI (stocks prepared in DMSO) for 72 hours. Protein extraction was performed using RIPA buffer. MYC1 and MYC2 tumors were homogenized in RIPA buffer. Dilutions of the antibodies were 1:1000 for S1PR1, S1PR2, pSTAT3 ser727, pERK, cMYC and GAPDH, and 1:5000 for secondary polyclonal anti-rabbit antibodies.

### RNA seq

*Sphk2* global KO mice and *Sphk2* WT mice were anesthetized approximately after 10-20 days of implantation. Tumor tissues and lymph nodes were harvested, homogenized with Kimble™ Kontes™ Pellet Pestle™ Cordless Motor homogenizer (DWK Life Sciences 7495400000) in RLT buffer and run through QIAshredder spin columns (QIAGEN). RNA was extracted using RNeasy Mini Kit (Qiagen) with DNase I treatment. Quality was confirmed with Fragment Analyzer (Advanced Analytical Technologies Inc.), and samples were sequenced using Illumina NextSeq500 at the Massachusetts Institute of Technology BioMicro Center. The raw data was mapped to the mouse Ensembl GRCm38 reference genome, gene read counts were generated and were normalized using STAR (Dobin *et al*., 2013), Bioconductor R package Genomic Alignments (Lawrence *et al*., 2013) and DESeq2 (Love, Huber and Anders, 2014), respectively in-house.

Differential gene expression was calculated using edgeR package (version 3.26.8). The differentially expressed genes with absolute fold change, logFc >2 and p-value <=0.01 were considered significant. GO and KEGG pathway enrichment analysis was performed using GOstats package with up to 1000 differentially expressed genes ranked by ascending p-value.

The complete list of differentially expressed genes and pathway enrichment analysis in *Sphk2* global KO versus *Sphk2* WT tumors and lymph nodes are in **Supplementary Tables 1, 2, 4 and 5.**

GSEA with hallmark, KEGG, reactome, GO and biocarta gene sets was performed using Broad Institute GSEA software (Subramanian *et al*., 2005) to systematically analyze the data. Pathways with the absolute value of normalized enrichment score (NES) >2 and FDR q-value <0.05 were considered significant. Complete list of GSEA pathways regulated upon *Sphk2* global KO versus *Sphk2* WT tumors and lymph nodes are in **Supplementary Tables 3 and 6**, respectively.

### Flow cytometry

Tumors, lymph nodes and spleen were harvested from mice post implantation and strained through 70um strainer. Peripheral blood was collected using cardiac puncture and serum extracted using PBMC fractionation. Spleen samples were lysed using ACK buffer. All samples were stained with fluorescently tagged primary antibodies. Details of the antibodies used are in the STAR table and the antibody panel in **Supplementary Table 7**. Samples were run using Fortessa X20 (BD Biosciences). Fixable Viability Dye eFluor™ 780 (eBioscience™, Cat# 65- 0865-14) was used to exclude dead cells and FSC-A versus -H plot was used to select for single cells. Data was analyzed using FlowJo. Percentages of total CD8^+^T, FOXP3^-^CD4^+^T, FOXP3^+^CD4^+^T, NK, Mφ, TAM, MG, DC like and monocyte like were calculated out of live CD45^+^ cells. Percentages of cytokine (TNFα^+^, IFNy+ and GrzB+) expressing CD8^+^T, NK or FOXP3^-^ CD4^+^T cells were calculated out of total CD8^+^T, NK, or FOXP3^-^CD4^+^T cells respectively. Percentage of effector memory markers (CD44highCD62L^-^) expressing CD8^+^T was calculated out of total CD8^+^T cells. Percentages of MHCII^+^CD86^+^ Mφ, TAMs and DC like were calculated out of total Mφ, TAMs and DC-like cells respectively. MHCII^+^ TAM MFI percentage was calculated out of total TAMs. Relative populations of different classes of immune cells, or their ratios from tumors from flow cytometry data, were Z-score transformed to allow for comparisons between different arms and populations. Heat maps were generated using the Seaborn 0.9.0 package in the Python language environment.

Populations were gated as: CD45^+^ live cells – CD45^+^ viability dye^-^, Mφ – CD45highGr1^-^CD11c^-^ CD11b^+^F4/80^+^, TAM – CD45highGr1^-^CD11c^-^CD11b^+^F4/80^+^, MG – CD45low/interGr1^-^CD11c^-^ CD11b^+^, DC like – CD45^+^Gr1^-^CD11c^+^CD11b^-^F4/80^-^, M1 like – MHCII^+^CD86^+^CD206^-^, M2 like – MHCII^-^CD86^-^CD206^+^, Monocytes – CD45^+^Gr1^-^ CD11c^-^CD11b^+^, CD8^+^T – CD45^+^CD8^+^, CD4^+^T – CD45^+^CD4^+^, Tregs or FOXP3^+^CD4^+^T – CD45^+^CD4^+^FOXP3^+^, FOXP3^-^CD4^+^T -

CD45^+^CD4^+^FOXP3^-^, Effector memory population – CD44highCD62L^-^, NK - CD45^+^CD3^-^CD19^-^ NK1.1^+^. Flow cytometry panel and gating are provided below in a table format.

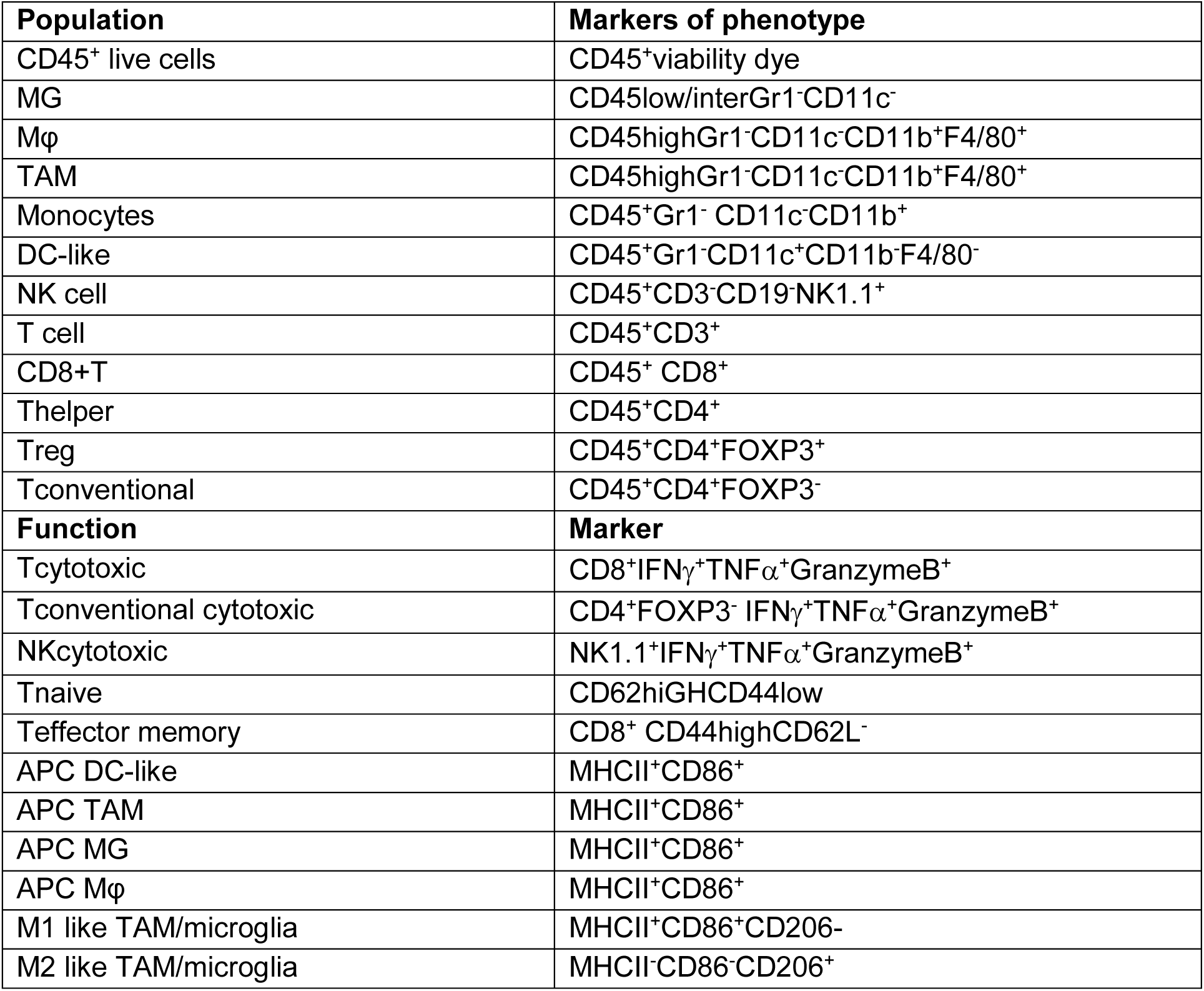

### FACS

Tumor samples were processed **(refer to flow cytometry section).** TAMs gated as live CD45highGr1^-^CD11c^-^CD11b^+^F4/80^+^ were sorted by BD FACSAria™ Fusion and collected in RIPA buffer.

### CD8 depletion studies

Depleting antibody targeting cell surface receptor CD8α was purchased from BioXCell (clone 53- 6.7, Cat# BE0004-1). Depleting antibodies were administered I.P. in mice on days 2, 4, 6, 8, 12, 16, 20, and 24 after tumor implantation. Animals were monitored for survival.

### Statistical Analysis

All data are expressed as mean ± SEM. Statistical analysis was performed in GraphPad prism. ANOVA one or two way was used to calculate significance. The Dunnett post doc test was performed after ANOVA to compare means to a control mean wherever applicable. Tukey-Kramer test was performed after ANOVA to compare every mean with every other mean while allowing for the possibility of unequal sample sizes wherever applicable. Two-tailed t tests were used between data comparing only two groups. Survival curves are plotted with the Kaplan-Meier method. For survival data, a log-rank test was employed. p<0.05 was considered significant across all types of analyses.

## Acknowledgments

We would like to acknowledge the support of Nour and Schindler families as patient advocates from Massachusetts General Hospital for this study. We would like to thank Drs. H.J. Kim, M. Pittet, G. Freeman, and T. Hla for invaluable suggestions, and Drs. G.B. Ferraro and S. Krishnan for helpful discussion. We would like to acknowledge Drs. M.F. Roussel, A. Garancher, and R. Wechsler-Reya for providing the MYC1 cell line and MYC2 primary cells as well as technical support with mouse model establishment and Drs. H. Willers and A. Luster for sharing experimental equipment. We would like to thank J. Fung and Dr. M. T. Chow for their expert technical assistance in flow cytometry, Dr. H. Lee for expert advice in statistical analysis. We would also like to acknowledge Drs. Peigen Huang, S. Yan, and G.A.V. Cruzeiro, as well as S. Roberge, T.H. Tale, C.J. Smith, A. Sanjali, and R.R. Ramjiawan for experimental assistance.

This study was supported by National Institutes of Health (NIH) grant R35CA197742 (to R.K.J). R.K.J also acknowledges support from the NIH (R01CA208205, R01-CA259253, R01-NS118929, U01-CA261842, U01CA224173 and U01-CA 224348), the Ludwig Center at Harvard, the Jane’s Trust Foundation, the National Foundation for Cancer Research and the Niles Albright Research Foundation. RNA-sequencing work was partially funded by Koch Institute Support (core) grant P30-CA14051 from the National Cancer Institute (NCI). S.C acknowledges support from the American Brain Association (ABTA Basic Research Fellowship), Massachusetts General hospital (MGH FMD Medical Discovery award), Pediatric Cancer research Foundation (PCRF Young Investigator’s Award), Jane Coffin Childs Memorial Fund for Medical Research (Jane Coffin Child Memorial Fellowship). P.J.L is supported by MGH ECOR FMD Fellowship. D.G.D acknowledges support from the NIH (R01CA274254 and R01CA260857) and a Katz Investigator Award. L.B and V.D acknowledge support from the Gates Foundation (OPP1140482). W.H.H received fellowship from the Agency for Science Technology and Research (A*STAR) in Singapore. A.S.K is supported by the Agency for Science Technology and Research A*STAR NSS (PhD) graduate fellowship. M.D is supported by the AACR-Loxo Oncology Pediatric Cancer Research Fellowship. Y.Z acknowledges support fromthe China Scholarship Council (File No.: 201406380077). N.P.T is supported by The CRI Irvington Postdoctoral Fellowship. P.A. was supported by the International Postdoctoral Fellowship from the Swedish Research Council.

## Author Contributions

Conceptualization, S.C. and R.K.J.; Methodology, S.C., P.K., Y.Z., H.-J.K.; Validation, S.C., P.K., Y.Z, S.K, and L.B.; Investigation, S.C., P.K., A.S.K, P.J.L, Y.Z, M.D., W.W.H, P.J.L., N.P.T, P.A, W.J.K. and S.J.W.; Formal Analysis, S.C., P.K., W.W.H, P.J.L. and A.S.K., Software, S.C., P.K., P.J.L., W.J.H. and A.S.K.; Resources, D.F, V.D. and R.K.J.; Data Curation, S.C., P.K., P.J.L., W.W.H., and A.S.K.; Writing – Original Draft, S.C. and P.K.; Writing – Review & Editing, S.C., P.K., S.K., A.S.K., W.W.H., N.P.T., P.A., W.J.K., S.J.W., D.G.D., D.F., D.H.E., T.I.Y., H.-J.K., and R.K.J.; Visualization, S.C., P.K., P.J.L., W.W.H. and A.S.K.; Funding Acquisition, S.C. and R.K.J.; Supervision, L.X. and R.K,J.

