## Supplementary figures and images for "Targeting Tumor-derived Sphingosine Kinase 2 Unleashes Antitumor Immunity and Improves Survival of Mice with Group 3 Medulloblastoma"

### Supplemental Figures

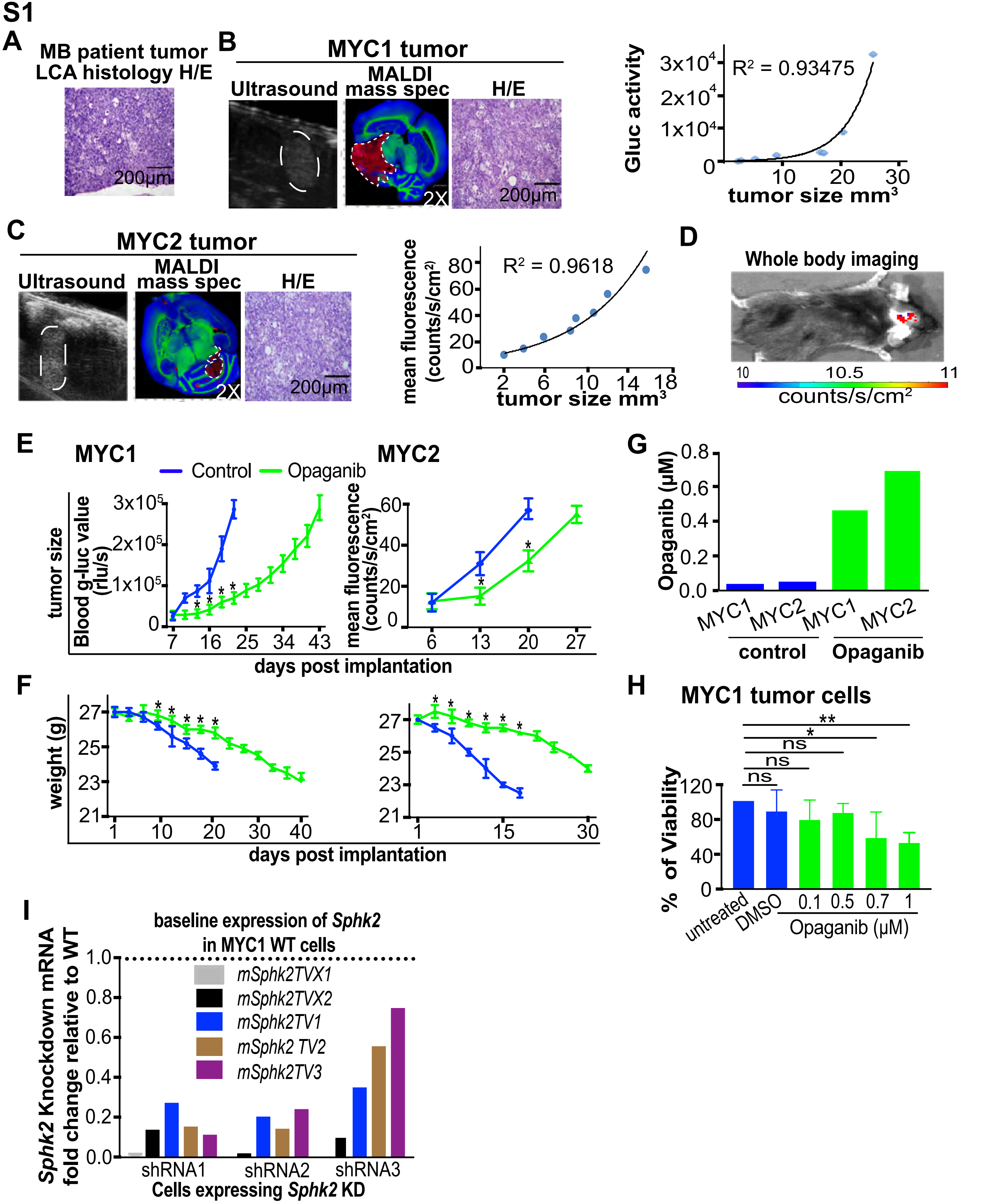

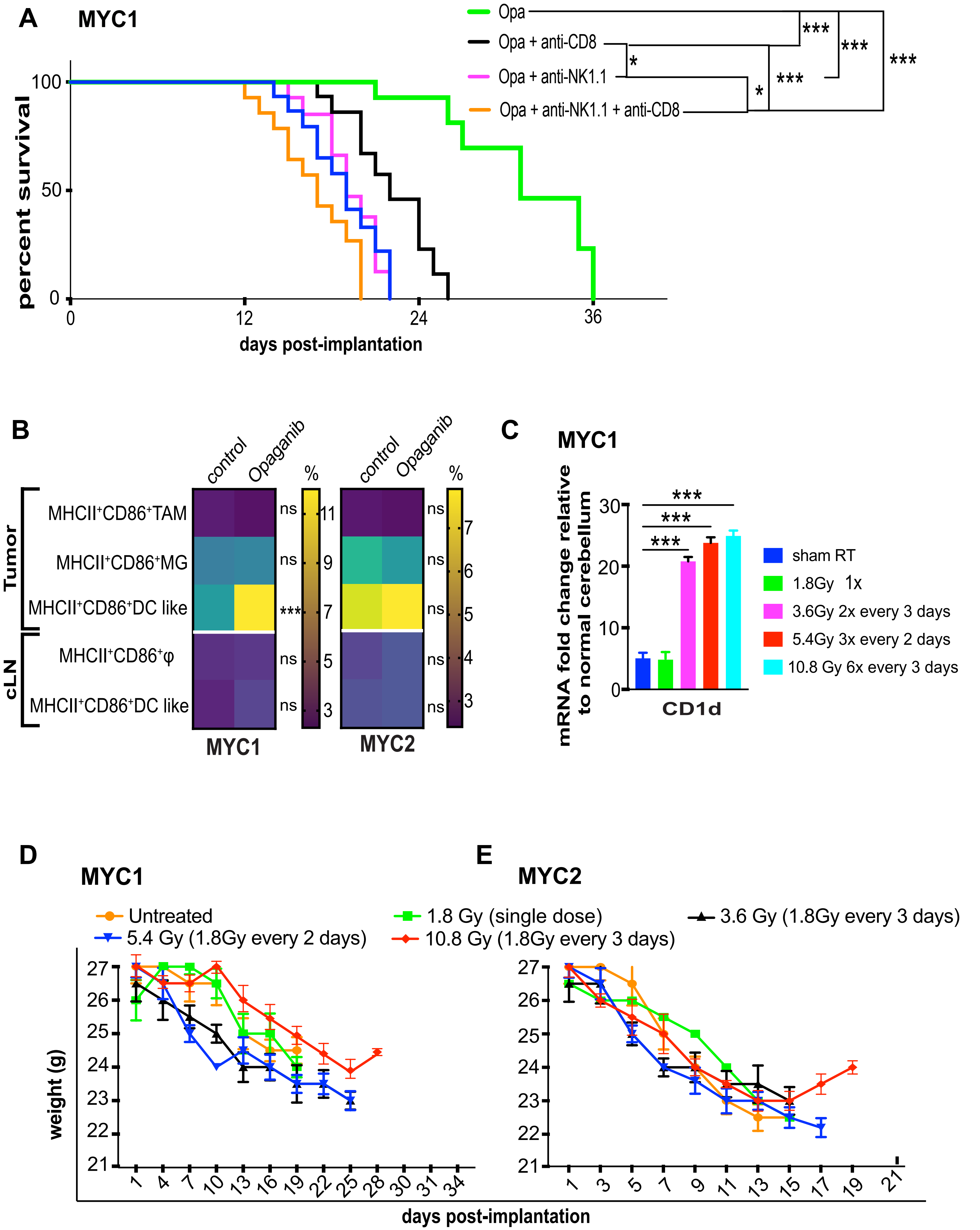

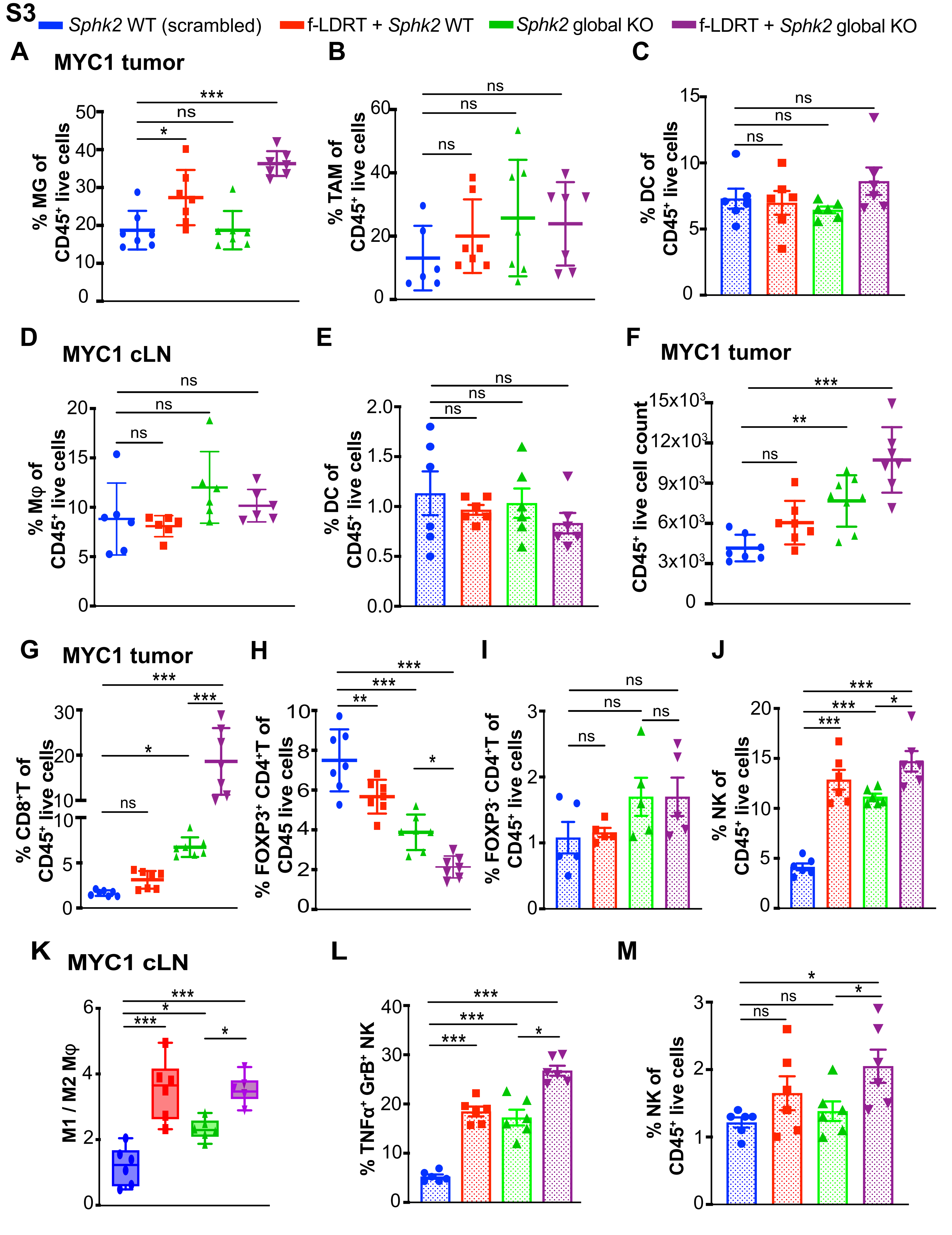

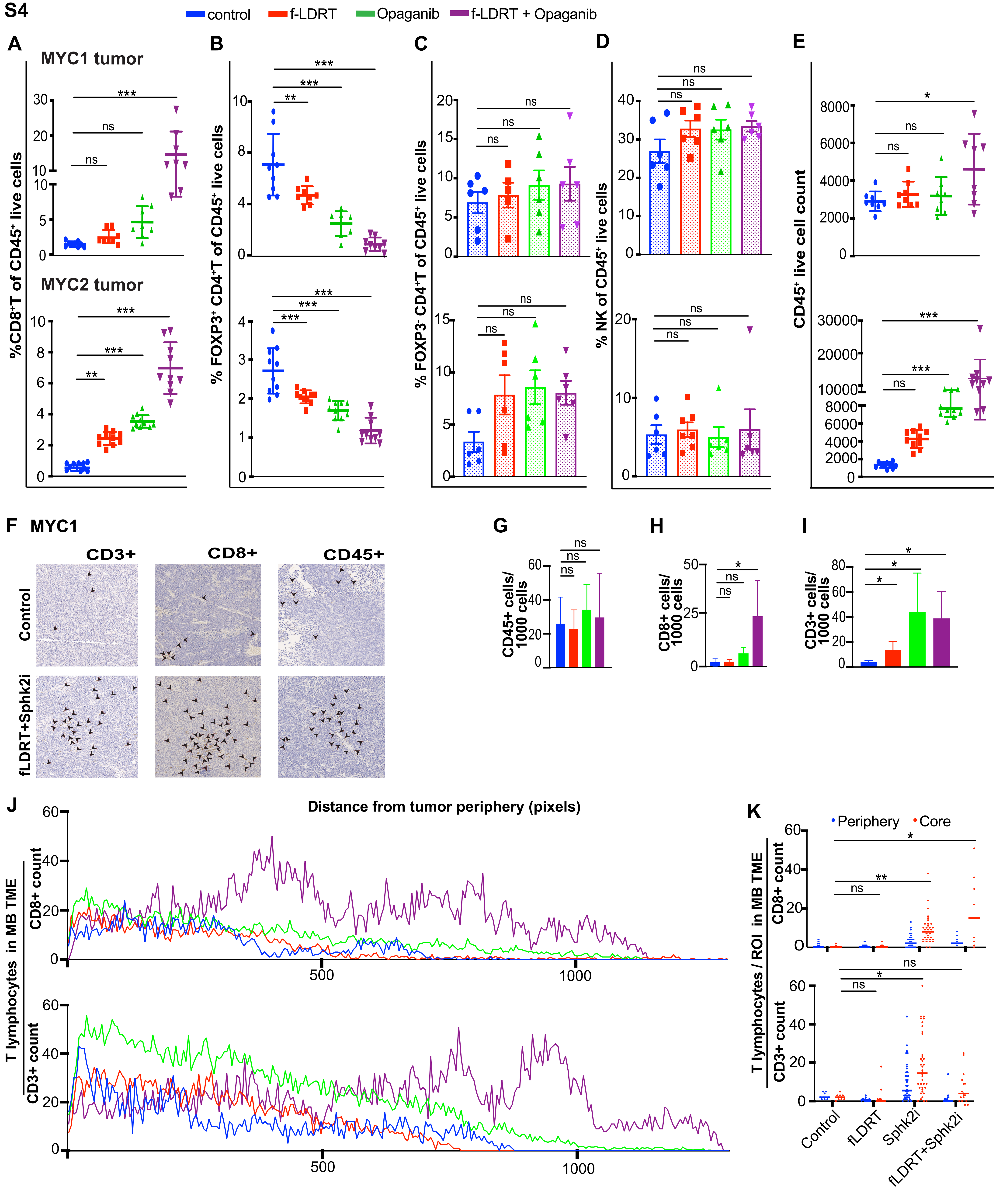

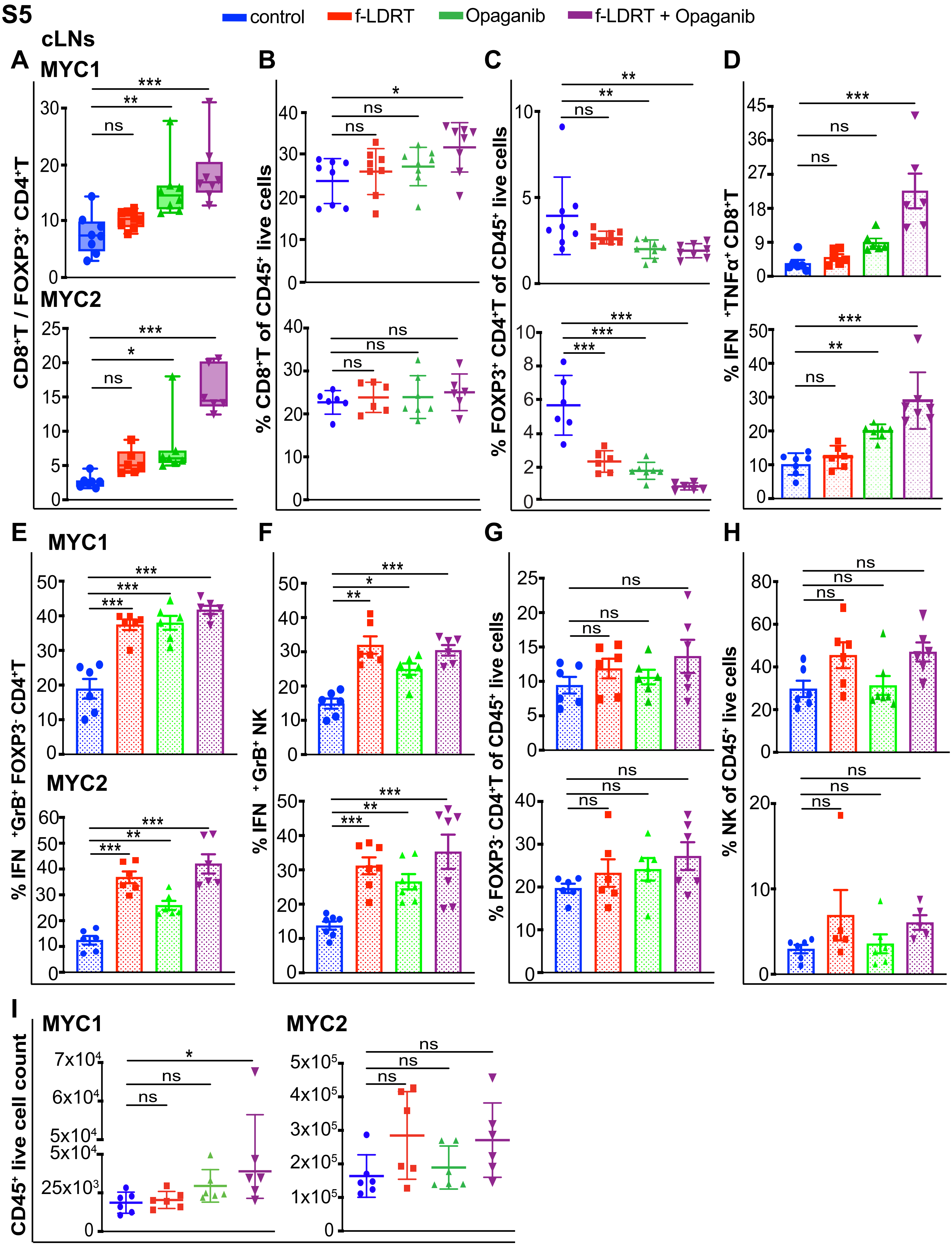

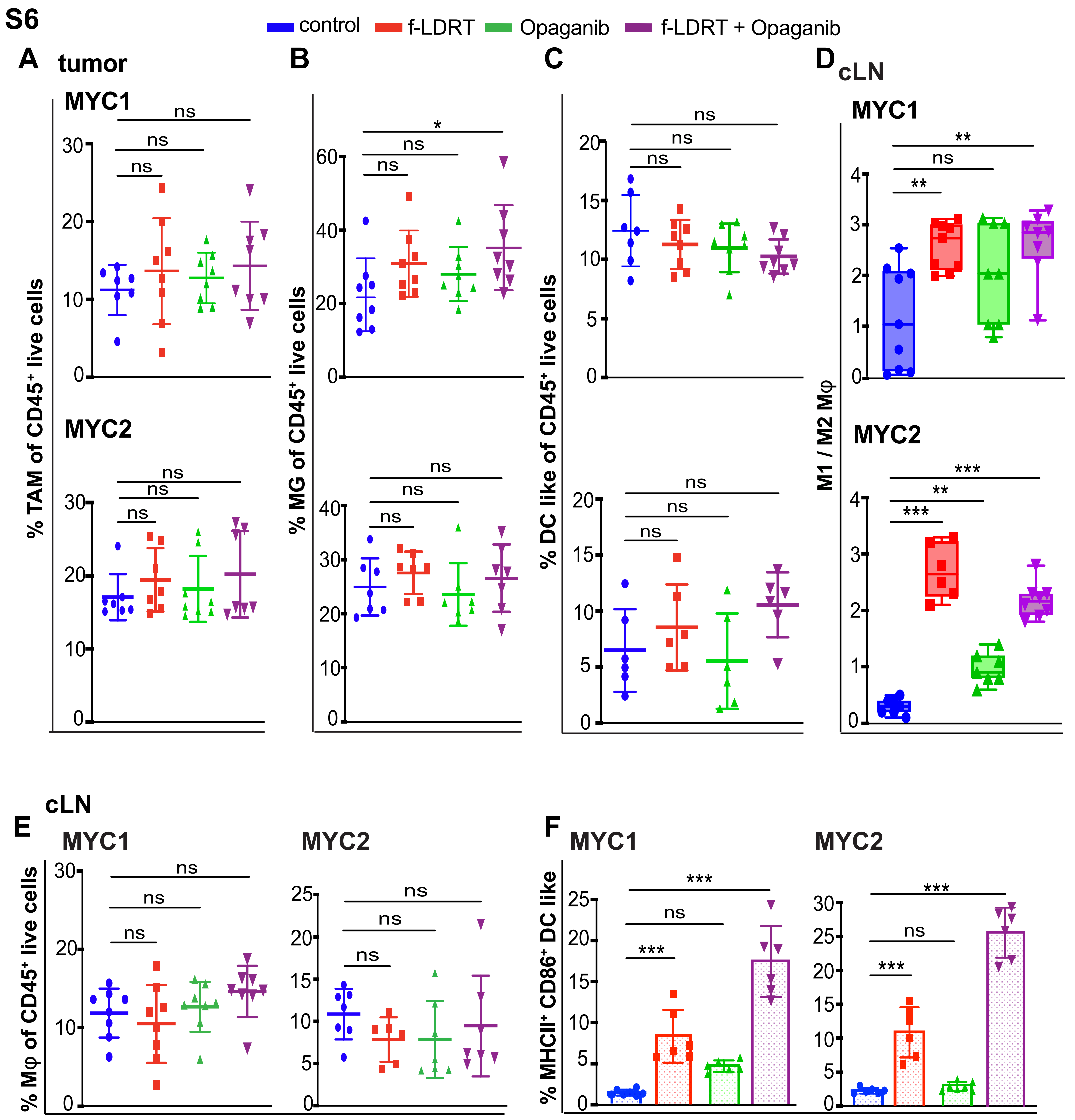

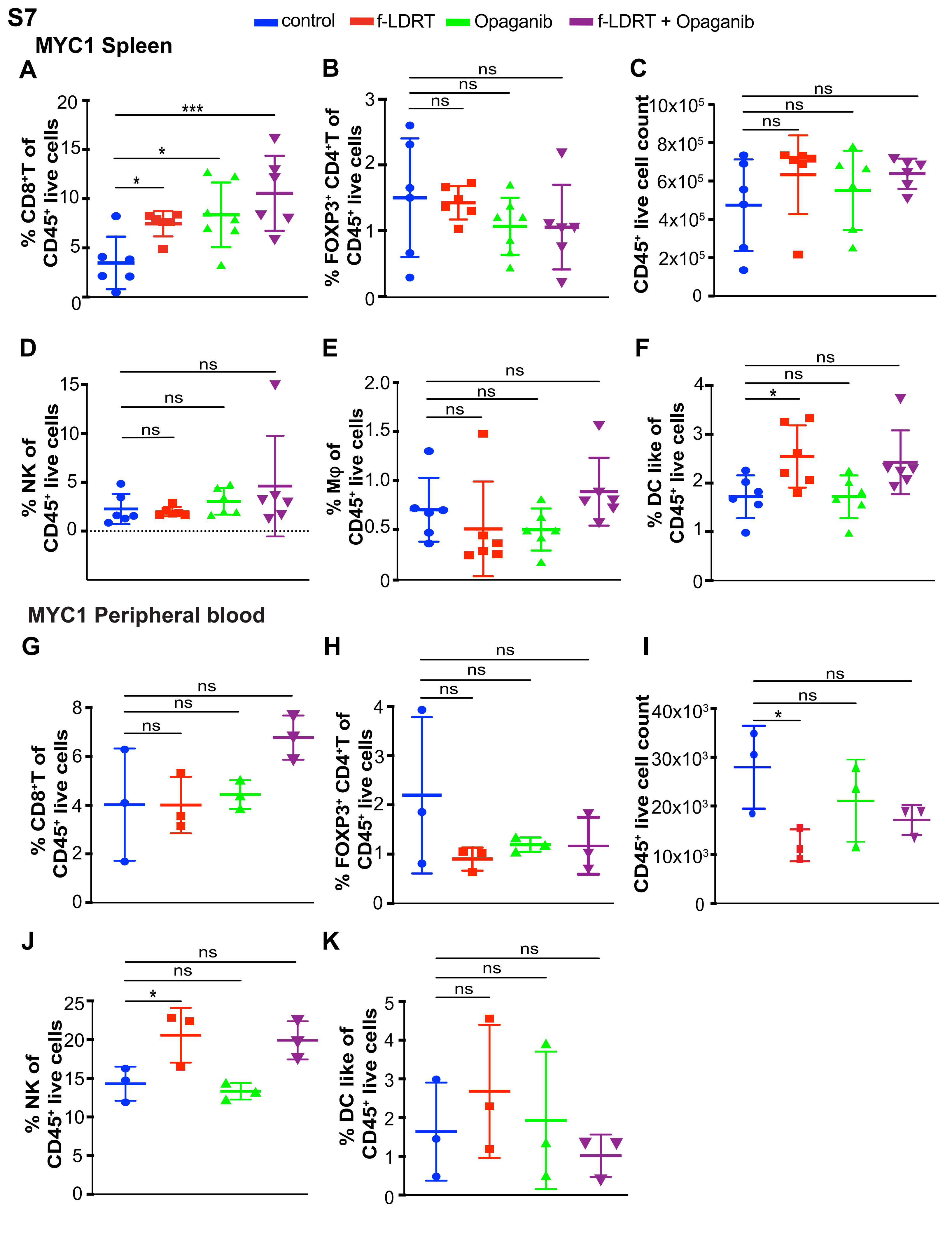
